# Metabolic Engineering on a 3D-Printed Microfluidic Platform: A New Approach for Modular Co-Metabolic pathways

**DOI:** 10.1101/2023.08.22.554264

**Authors:** Seyed Hossein Helalat, Islam Seder, Rodrigo C. Téllez, Mahmood Amani, Yi Sun

## Abstract

Metabolic engineering of cell factories often requires extensive modification of host cellular machinery, leading to numerous challenges such as metabolic burden, intermediate metabolite toxicity, and inadequate endogenous fluxes. To overcome the limitations, we presented an innovative approach for metabolic engineering, by constructing modular biosynthetic pathways on a 3D-printed microfluidic platform. Several new techniques have been developed, including novel designs of chip configurations, effective methods for enzyme immobilization on printed resins, and proper ways to regenerate cofactors in redox reactions. As a proof of concept, we built xylose consumption and CO_2_ fixation pathways in the microfluidic chips and successfully demonstrated that the platform was able to convert xylose and enable the rapid growth of *Saccharomyces cerevisiae,* which otherwise will not grow with xylose as the only carbon source. Overall, the 3D-printed microfluidic platform presents a much simpler and more efficient cell-free strategy for developing modular, optimized biosynthetic pathways.

## Introduction

Microbial cell factories have been increasingly employed for the bioconversion of substrates into valuable products^1^. Biosynthetic production can utilize a wide range of feedstock, proceed under physiological conditions, and avoid environmentally deleterious byproducts, thus presenting a sustainable alternative to the prevailing chemical conversion processes^2^. In order to produce diverse metabolites, researchers in synthetic biology and metabolic engineering have endeavored to extensively modify the host cellular metabolic machinery by constructing novel pathways and optimizing metabolic fluxes^3^. However, they have encountered several challenges, such as low efficiency, elevated metabolic burden, accumulation of toxic intermediates, substrate diffusion limitations, and inadequate endogenous metabolic flux toward desired target products^4,5^. Additionally, integrating heterologous enzymes from other microorganisms into producer hosts is problematic due to their requirement for different conditions in terms of temperature, cofactor concentrations, and ionic conditions^6,7^.

One strategy to overcome these obstacles is modular co-culture metabolic engineering. It divides a complete biosynthetic pathway into separate serial modules, with individual modules distributed across different microbial strains to express associated genes and attain the desired biosynthesis. This method has shown improved product titers that surpassed monoculture. However, limitations associated with engineering and culturing multiple host modules remain apparent^8–10^.

Alternatively, there has been great interest in conducting bioconversion using *in vitro* cell-free systems, which couple various enzymes for certain metabolic pathways and create enzyme cascade reactions to simulate the multi-step metabolic flow in real cells^11^. These biomimetic systems allow for the flexible combination of enzymes from different hosts; meanwhile, the enzymatic reaction conditions can also be fine-tuned to achieve optimal efficiency. The cell-free approach thus could potentially remove the constrains of living organisms and provide a simpler and more streamlined tool to build metabolic pathways that are missing or inefficient in natural/engineered cell factories^12,13^.

There are various ways to construct multienzyme bioreactors. One type of structures is called enzyme microcarriers, where the selected enzymes are sequestered in separate microcompartments made of *e.g.* oil-water droplets^14^, hydrogel microcapsules^15^, liposomes or polymersomes^16^. These microcarriers are relatively simple to generate, whereas it has been difficult to control the arrangement of the enzymes as well as the intermediate flow between the compartments, resulting in low catalytical efficiency. Alternatively, microfluidic chips fabricated in glass, Polydimethylsiloxane (PDMS) or plastics have been developed as promising cascade enzyme reactors^17^. Improved control has been achieved in microfluidic devices since the enzymes are immobilized on the chips, and the substrate solutions are transported by external forces to react with the enzymes for continuous biocatalysis. However, a few issues are hindering their translation from laboratory to industrial production, such as the high cost of microfabrication, insufficient mixing and reaction kinetics, lack of cofactor regeneration, poor enzyme immobilization, *etc.* All these factors may negatively impact the final metabolite titer, production rate, and yield, making the enzymatic microreactors far from real biosynthesis applications^17^.

Recently, three-dimensional (3D) printing technology has emerged as a new microfabrication technology and attracted considerable interest. By harnessing the capabilities of high-resolution 3D printers, sophisticated 3D microfluidic devices can be rapidly fabricated in a single step from a computer model^18^. The advantages such as flexible design, fast prototyping, facile upscaling, cost-effectiveness, and availability of biocompatible resins, open up new doors to develop industrial-applicable microfluidic systems for multi-enzymatic reactions^18,19^. However, to our best knowledge, no work has been devoted to develop an integrated 3D-printed platform that can realize complex metabolic pathways.

In this study, we aimed to devise 3D printed microfluidic platforms dedicated to metabolic engineering on chip (**Fig. 1**). The successful implementation of this approach needs a number of new techniques and methods, such as the new design of chip configurations, effective methods for enzyme immobilization on printed resins, and proper ways to transfer electrons to the enzymes in redox reactions. To achieve efficient mixing and heat transfer for catalytic reactions, we designed a microfluidic chip with a snake-like channel configuration in conjunction with irregular barrels, and fabricated the chip by stereolithographic 3D printing using a biocompatible resin. Then, we managed to establish a chemical method to immobilize enzymes on the 3D-printing resin surface with high bioactivity. We showed that the amount of immobilized enzymes could be significantly augmented by coating dendrimer nanoparticles on the channel surface. Moreover, we innovatively inserted a screen-printed electrode into the microfluidic chamber to facilitate redox reactions, in which efficient electron transfer to the enzymes was achieved through direct electron transfer and co-factor reduction approaches.

**Figure 1.**
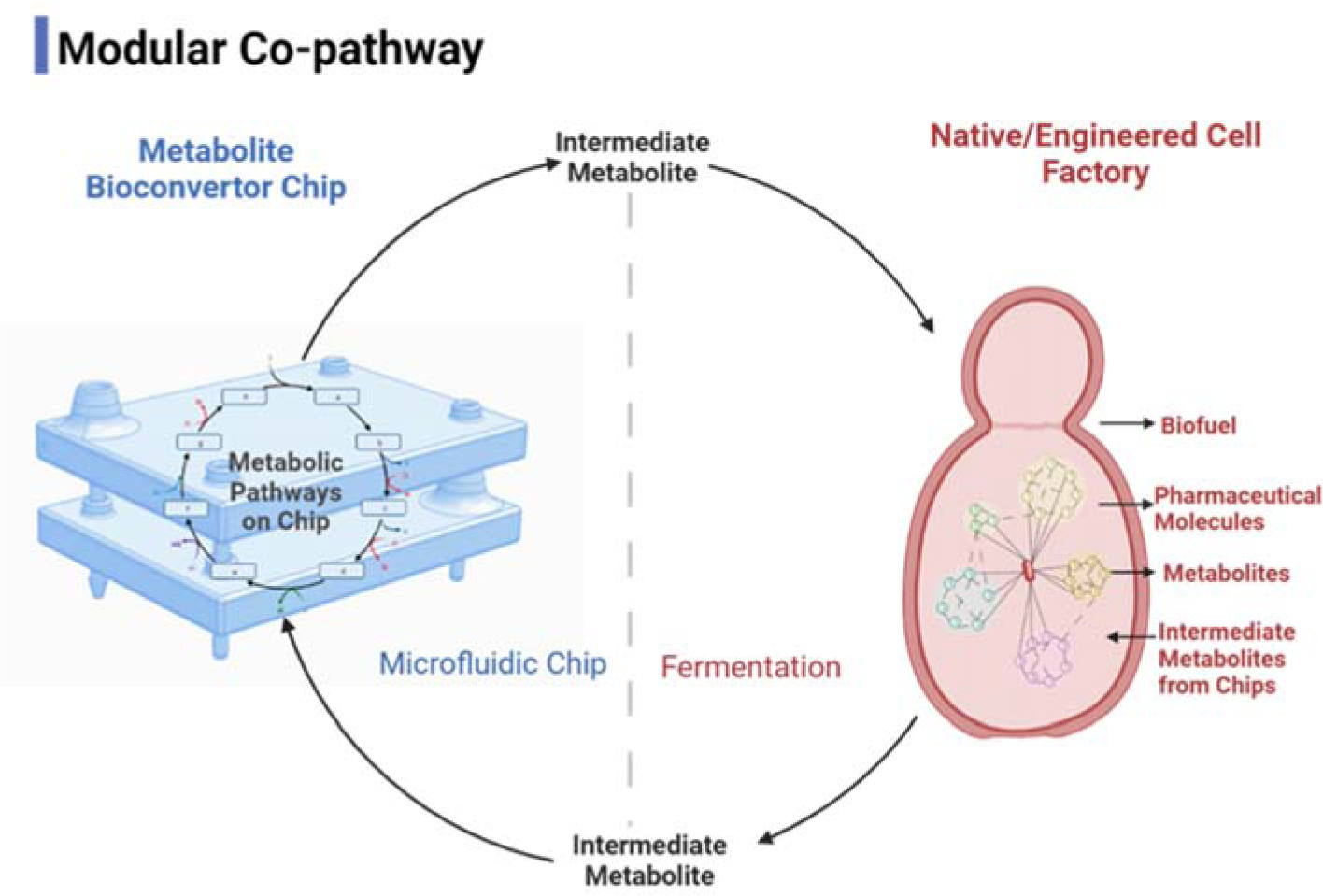
A novel cell-free modular co-pathway system on a 3D-printed microfluidic platform, aiming at augmenting metabolite production. This system allows the seamless interconnection of diverse engineered cells and chips, forming a tandem conversion system. The application scenarios are versatile, for instance, 1) the integrated pathway within the chip can serve as a carbon source supplier for native or engineered cells; 2) an intermediate metabolite derived from the chip can undergo conversion by the cell, leading to the desired product; 3) intermediate metabolites synthesized from simple carbon sources by the cell can be subsequently transformed into the final product by the chip.

To demonstrate the potential of the 3D-printed microfluidic platform for metabolic engineering, we have constructed two modular co-metabolic pathways consisting of six enzymes to enable *Saccharomyces cerevisiae* to grow with xylose as the carbon source. We immobilized the six enzymes on three chips and inserted the electrode in one of the chips to regenerate the cofactor for the redox reactions. The yeast growth characterizations revealed that the 3D-printed microfluidic platform successfully fermented xylose and assimilated CO_2_, allowing natural *S. cerevisiae* to sustainably produce diverse biofuels and chemicals from xylose while achieving a virtually carbon-neutral fermentation process.

Overall, our work showed that the 3D-printed microfluidic platform presents a very facile and efficient strategy to develop modular metabolic pathways, thereby circumventing the enormous effort needed to engineer the host cells. We envision the new methodology would hold great promise in biotechnology, metabolic engineering, and sustainable fermentations.

## Material and Methods

### Microfluidic chip design and simulation

SOLIDWORKS 2021 software (Dassault Systems) was utilized for designing the enzymatic reaction chips and simulating the fluid flow dynamics within the channels. Various chip shapes, including Y, circle-end diamond, two-square, bubbly, snake, arranged-column, skewed-column, and Tesla-design, were investigated. The influence of barriers on enhancing mixing within the channels was also studied, followed by adjustments in barrier size and location.

### Chip fabrication and experimental setup

The integrated 3D-printed microfluidic platform was assembled using a series of enzymatic reaction chips, an electrode, a gas chamber, and a cell growth chamber. All of these chips and chambers were fabricated utilizing 3D printing technology and were constructed from a biocompatible resin (BioMid resin, Formlabs), employing a digital light processing (DLP) 3D printer (Asiga Pico2 HD, Asiga). Consequently, after optimizing the printing conditions, the printer settings were established with an exposure time of 5 seconds and a resolution of 50 μm. After the printing process, the chips underwent a thorough rinsing with isopropyl alcohol (IPA) for a duration of 20 min, followed by a 15-min sonication step and a wash with IPA. Then, the chips were subjected to a 2-h exposure to UV light at 65 °C to ensure crosslinking of the uncured resin. Further treatment involved an overnight incubation at 80 °C, followed by washing with 20% ethanol and sterilized water.

For the cell chamber chip, a 0.22 μm pore size membrane (47 mm-Manga Nylon Filter, GVS-Lifescience) was inserted in the middle of the chamber to separate the cells from the circulating fluid medium. When the redox reaction was involved, an electrode (C220AT Screen-Printed Gold, Metrohm) was inserted into the reaction chip and secured with resin to prevent leakage between the chamber’s side wall and the electrode. The electrode was connected to a potentiostat (Autolab, Metrohm) to get the voltage supply. The chips were interconnected and connected to the micropump (Mp-advance kit, Bartels Mikrotechnik) via tubes (Tube Tygon LMT-55, Saint-Gobain). During operation, the chips were heated to 37 °C using the provided heater to maintain the desired temperature. The flow rate through the pump was controlled using software (Multiboard, Bartels Mikrotechnik). In order to facilitate sample intake and retrieval, a special inlet and outlet were incorporated into the cell chip, allowing for convenient handling with a pipette.

### Chip activation and enzyme immobilization

To immobilize the enzymes within the inner channels of the 3D-printed microfluidic chips, a process involving the crosslinking of 3-aminopropyl trimethoxysilane (APTS) and glutaraldehyde was used. This technique facilitates interaction between the amine groups present on the protein’s surface and the crosslinking agents. Following the curing of the chips, the channels underwent a series of washing steps with distilled water and were subsequently dried using nitrogen gas. Afterwards, a pre-warmed (55°C) solution comprising ammonium hydroxide and hydrogen peroxide (in a ratio of 3:1) was introduced into the channels, followed by a 5-min incubation at the same temperature. After removing the solution, the channels were washed three times with distilled water. Next, a solution of APTS and water (at a ratio of 1:9) was introduced into the channels and allowed to incubate at room temperature for 2 h. The solution was then removed, and the channels were subjected to five washes with distilled water.

In the next step, a solution consisting of 25% glutaraldehyde and water (at a ratio of 1:4) was added to the channels and incubated at room temperature for 1 h. After incubation, the solution was removed, and the channels were washed five times with 20 mM Tris-HCl buffer. Finally, the protein of interest, suspended in 20 mM Tris-HCl buffer at a concentration of approximately 1-5 mg/mL, was introduced into the chip, which was subsequently kept at 4°C overnight. On the following day, the solution containing unbound proteins was removed, and after undergoing ten washes of the channels with 20 mM Tris-HCl buffer, the chip was ready for use.

To further enhance the amount of immobilized enzymes within the channels and improve the efficiency of enzymatic reactions, polyamidoamine (PAMAM) dendrimer nanoparticles (generation 5.0) from Sigma were utilized for coating the channels. Following the APTS and glutaraldehyde steps in the chip activation process, dendrimers at a concentration of 25% (v/v) were introduced and incubated for 3 h at room temperature. Afterwards, the unbound dendrimers were removed, and a thorough wash with distilled water was performed. Next, a solution of 25% glutaraldehyde and water (at a ratio of 1:4) was added to the channels and incubated at room temperature for 1 h. The channels were then washed five times with 20 mM Tris-HCl buffer. Finally, the proteins of interest were introduced and allowed to incubate overnight. On the subsequent day, unbound proteins were removed, and the channels underwent ten washes with 20 mM Tris-HCl buffer to ensure proper preparation for subsequent experiments.

### Enzymatic reaction assay on the chips

In order to investigate the chip design and evaluate the effectiveness of chemical modifications, the enzymes lactate dehydrogenase (LDH) and β-galactosidase from Sigma were conjugated to the microfluidic chips, and the enzymatic reactions were examined under various experimental conditions. The conversion of pyruvate to lactate by lactate dehydrogenase was assessed with a pyruvate assay kit (Sigma, MAK332), and the color change of X-gal was observed in the channels and buffers for the β-galactosidase assay.

### Electron transfer to the enzymes

Two strategies were examined to facilitate electron transfer: direct electron transfer and indirect electron transfer with co-factor NAD(P)^+^ reduction/oxidation. The LDH was used as a model enzyme. In the case of direct electron transfer assessment, the lactate dehydrogenase enzyme was immobilized directly onto the surface of electrodes (C220AT Screen-Printed Gold from Metrohm), obviating the need for NAD(H) in the reaction buffer. Only 150 mM pyruvate was added to the reaction buffer as the substrate. For the co-factor (NAD+) reduction/oxidation strategy, immobilization of lactate dehydrogenase enzyme was accomplished in a 3D-printed chip with the electrode inserted to the chip. 150 mM pyruvate and 15 mM oxidized form of the co-factor (NAD^+^) were added to the reaction mixture. A range of voltages (-(0.1–9)) in varied concentrations of PBS (0.1-1 dilutions) and Tris-HCl (20 mM) buffers were assessed to optimize electron transfer efficiency. The conversion of pyruvate to lactate was quantified using a pyruvate assay kit (Sigma) to evaluate the effectiveness of these strategies. Moreover, as LDH is capable of converting lactate into pyruvate, we conducted experiments to assess the production of pyruvate from lactate. To achieve this, we reversed the reaction by adding 150 mM lactate and 15 mM NADH to the reaction buffer, while applying positive voltages.

### Computational pathway design

The integration of the xylose consumption pathway with the CO_2_ fixation pathway from *E. coli* was achieved using the Cytoscape StringApp software. Distinct protein networks were constructed for xylose conversion, glycolysis, pentose-phosphate, and CO_2_-related proteins with confidence (score) cutoff of 0.4. Initially, a comprehensive list of proteins was extracted from UniProt, PubMed query, and universal interaction databases specifically pertaining to CO_2_-related processes. Subsequently, these networks were merged to establish connections between the various protein components. Then, the most closely associated proteins, possessing the capability for CO_2_ fixation, to xylose consumption pathway within these networks were identified and characterized.

### Strains and Enzyme production

To facilitate plasmid propagation, we employed *E. coli* NM522 strain, which was cultured in LB medium (10 g/L tryptone, 5 g/L NaCl, and 5 g/L yeast extract) at 37 °C with continuous shaking at 250 rpm. To establish the xylose consumption and CO_2_ assimilation pathway, we amplified the *xylA* (UniProt: P00944), *xylB* (UniProt: P09099), *rpe* (UniProt: P0AG07), *gnd* (UniProt: P00350), *edd* (UniProt: P0ADF6), and *eda* (UniProt: P0A955) genes using specific primers (Supplementary **Table S1**). These genes were then cloned into the pETDuet™-1 vector using the Gibson assembly kit. The resulting constructs were verified through sequencing, followed by transformation into the *E. coli* BL21 (DE3) strain for gene expression.

For protein expression, 2 mL of the overnight culture was transferred to 200 mL of fresh media containing 100 µg/mL ampicillin and incubated at 37°C (250 rpm) for 3 h. Once the OD_600_ reached approximately 0.8, IPTG was added to a final concentration of 0.5 mM to induce protein expression. The cells were then incubated overnight at 30°C to allow for optimal protein production. For protein purification, pellets from the 200 mL cultures were obtained, re-suspended in PBS, and lysed *via* sonication. Lysate was centrifuged and approximately 6 mL of supernatant was used for protein purification employing 200 µL of HisPur™ Ni-NTA Resin and Pierce™ Spin Columns (Thermo Fischer Scientific), following the manufacturer’s instructions. Of note, 40 mM imidazole was used as the washing buffer, according to in-house optimization tests. After purification, protein yield and integrity were assessed by analyzing 15 µL of the eluate using SDS-PAGE. To increase enzymes’ purity, they were further refined using Superose 6 (10/300) GL columns. Before use, the buffer of the proteins was exchanged, and the concentrations were adjusted using Amicon® Ultra-15 Centrifugal Filter Units (Merck) with the appropriate molecular weight cut-off, employing 20 mM Tris-HCl as the using buffer. The obtained purified proteins were stored at −80°C in 1 mL aliquots in a PBS solution containing 25% glycerol, 0.5 mM DTT, and 5 mM MgCl_2_ to maintain their stability.

### Enzymatic reaction conditions

Initially, the enzymatic reactions were carried out in tubes. A Tris-HCl 20 mM buffer supplemented with 5 mM MgCl_2_ and 0.5 mM MnCl_2_ was utilized as the fundamental buffer solution. A mixture consisting of 10 mM xylose, ATP, and 20 mM NADPH, along with 1 mM MgCl_2_, was added to the buffer solution. Various combinations of enzymes were investigated to study the individual reactions within the pathway. Specifically, XylA was employed as the starting point to catalyze the conversion of xylose, followed by the stepwise addition of other enzymes to complete the pathway. The reaction temperature was maintained at 37°C throughout the experimental procedure. The reaction products obtained after 3, 12, and 24 h were subjected to HPLC analysis and yeast growth characterization.

The enzymatic reactions were then evaluated on microfluidic chips to investigate the pathway with the immobilized enzymes. Upon applying the electrode to the microfluidic chip, the NADPH concentration was reduced by a factor of 20 to 1 mM, and a constant voltage of −1V was supplied to the electrode. Concerning the enzymatic reactions involving the Gnd enzyme in tubes, we conducted these reactions in an environment containing 20% CO_2_ to enhance the enzyme’s carboxylation capability. Additionally, within the chip system, we introduced a CO_2_ gas chamber to augment the concentration of CO_2_ during the process. Moreover, the stability of the immobilized enzymes in chips was compared to the free enzymes in tubes over a span of 7 days.

### Yeast culture conditions

A minimal medium devoid of sugar and peptone/tryptone carbon sources was prepared. To assess the yeast’s capacity to metabolize intermediate metabolites derived from the engineered pathway, we cultivated the yeast in the minimal medium supplemented individually with the pathway-specific intermediate metabolites, including xylose, xylulose, xylulose 5-phosphate (Xu5P), ribulose 5-phosphate (Ru5P), 3-deoxy-2-keto-6-phosphogluconic acid (KDPGA), 6-phosphogluconic acid (6-PGA), pyruvate, and glyceraldehyde 5-phosphate (GA3P). The yeast was incubated in a flask at 30 °C with continuous shaking at 200 rpm. The growth of the yeast was monitored at 24, 48, and 72-h time points using spectrophotometric analysis at an absorbance of 620 nm. Since we added ATP and NADPH to the reaction buffer to support the enzymatic reactions within the pathway, ATP, ADP, NADPH, and NADP^+^ were tested for their influence on yeast growth. To evaluate the growth of the yeast in the presence of pathway metabolites, enzymatic products obtained from the in-tube or on-chip reactions were supplemented to the minimal medium, and the growth of the yeast under different conditions was assessed.

Besides culturing the yeast in conventional flasks, we also fabricated the yeast growth chamber by 3D-printers and connected it to the microfluidic chip system. Two distinct strategies, namely batch-to-batch connection and continuous connection, were examined. In the batch-to-batch system, the reactions were initiated within the microfluidic chips, and after 24 h, the valve connecting the yeast growth chamber was opened. This approach allowed the periodic addition of substrate to the reaction every 24 h. Conversely, in the continuous system, the microfluidic chip and the yeast growth chamber were initially interconnected, facilitating simultaneous operation. Both systems were rigorously evaluated, and the growth performance of the yeast was measured and analyzed.

### HPLC

High-performance liquid chromatography (HPLC) (Shimadzu Co.) with an ion exchange column (Agilent Hi-Plex Column) was employed to detect the metabolites of first part of pathway and the final product. Xylose, xylulose, Xu5P, ADP, ATP, NADPH, NADP^+^, and pyruvate (Sigma Co.) were used as the standard samples to evaluate the retention time. Regarding the running conditions, 20 µL of samples were injected, the flow was set to 0.6 mL/min, and the temperature for the column and detector were at 60 °C and 55 °C, respectively. 5 mM H_2_SO_4_ was used as mobile phase, and RI and UV (190-700 nm) detectors were used to recognize the molecules. Samples from the chip, tubes, and yeast fermentation were filtered before injection with syringe filters (0.22 µM) and the Amicon column (3 KD) to remove the proteins and residuals.

## Results

### Chip design

In a microfluidic reactor, since the enzymes are immobilized on the chip, the reaction performance is largely influenced by the accessibility of the substrate to the immobilized enzymes. Consequently, the configuration of the chip has a substantial impact on the overall biocatalytic reaction. A few features are desired for the chip to achieve the highest possible reaction kinetics, such as efficient mixing to enhance mass transport, large surface area to increase the amount of the enzyme, as well as small pressure drop to minimize the damage to the enzyme activity^17^. Our primary objective was to devise and manufacture a novel channel configuration specifically tailored for this purpose. By utilizing computational simulations, we examined the flow dynamics of several channel designs, including snake-like, two-square, arranged-column, skewed-column, Tesla^20^, and snake-like with barrels (Supplementary **Fig. S1**).

Fluid flow behavior in the channels was evaluated by Reynolds number, Re = QD/νA, where Q and ν are the flow rate and kinematic viscosity of the fluid, and D and A are the diameter and cross-sectional area of the outer tube, respectively. We determined that the snake-like channel configuration, in conjunction with irregular barrels (TurboMix), exhibited superior mixing efficiency (**Fig. 2**). The TurboMix channel was specifically engineered to induce vortexes and whirlpools in the x, y, and z directions within the channels. We designed the snake-like channel in a relatively big size (ɸ = 2 mm, total length = 200 mm) to have a large volume (500 µL) and low pressure drop with an inner surface area of a plain chip as 1250 mm^2^. With the snake-like channel alone, Re is ∼10 at a flow rate of ∼1000 µL/min, imposing a laminar flow dominance inside the channel, making it difficult to have a vortex that assures efficient contact with the surface of the channel. Thus, we added microstructures inside the channel that create vortices to improve the mixing (**Fig. 2a-c**). The introduction of a column barrier leads to an alteration in the flow dynamics (Supplementary **Fig. S2a, b**). However, this also caused fluid stagnation (flow rate ∼0) behind the barrier (Supplementary **Fig. S2c**). To address this issue, we adjusted the positioning and increased the density of the barriers to ensure improved mixing throughout the channel (Supplementary **Fig. S2d**). The optimized TurboMix design chip and detailed dimensions are shown in **Fig 2d** and Supplementary **Fig. S3.**

**Figure 2.**
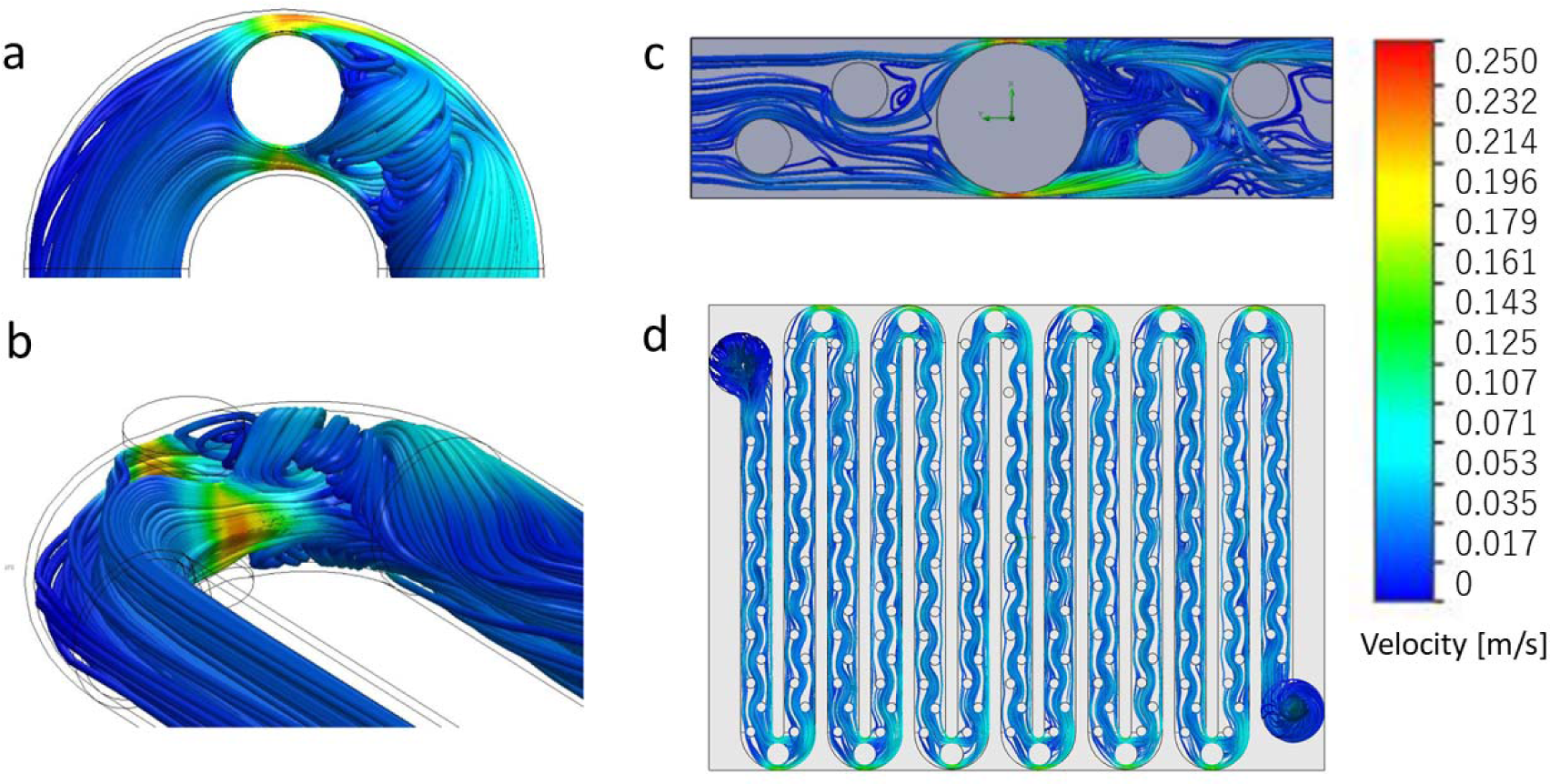
**a**-**c.** Simulations of effect of barriers on the flow in channels. **d.** Simulation of flow in TurboMix chip.

To examine the mixing, we let two solutions to flow in parallel inside the channel and characterize the mixing index at the outlet of the chip. The mixing index exceeds 90% for flow rate above 500 µL/min (Supplementary **Fig. S4**). Details of the mixing calculation and experimental set-up are included in Supplementary **Fig. S4**. Unless specified, we used flow rate of 500 µL/min. Moreover, within the TurboMix channel, the irregularly placed barrels also increase the surface area to 1720 mm^2^, resulting in a high surface-to-volume ratio of 3.4 mm-1, thereby enhancing the contact of the substrate to the immobilized enzymes. In addition, attributed to the relatively large dimension of the channel, the pressure drop was calculated to be 3 kPa. The small fluid resistance and pressure drop are favorable, as it ensures the enzyme activity is less affected.

### Chip manufacturing

Materials and fabrication methods of the metabolic chips play a crucial role in successful industrial translation. PDMS material is traditionally used at the research level for its ability to fabricate complex designs using soft lithography process. However, PDMS is a permeable material, thus fluid/media evaporates during a lengthy process. Importantly, PDMS chips need a multilayer chip design to achieve the desirable complex feature, and it requires several steps for fabrication, making it an unattractive choice for commercial manufacturing^21^. Instead, our metabolic system used 3D-printing technology with a biocompatible material (resin) to allow for facile scaling up, rapid fabrication, where the whole chip consisting of the channels, vortex generator, inlet, outlet, and support can be finished in a single process, which is not achievable by other mass production methods such as injection and extrusion molding.

Given the extensive array of resins available for 3D printing, our research necessitated the identification of a biocompatible silicon-based resin that possesses the capacity for high-resolution printing in order to achieve the intricate chip designs. Several resins (Grey Pro, Greyscale and BioMid resins from formlabs) were printed and tested using stereolithography (SLA)-based 3D printer. BioMid resin showed outstanding molding flexibility. However, SLA-based 3D printer failed to fabricate closed channels and small barriers (<2mm), thus, we shifted to the DLP-based printing (Asiga) with employing SLA-based bioMed resin. This combination of a suitable biocompatible resin and high-resolution printer (25 µm) achieved sophisticated metabolic platform (**Fig. 3a**). It only took 30 min to print a single chip. Examination of the printed chips showed that the patterns and dimensions were well aligned with the design, and no distortion or shrinkage was observed (**Fig. 3b**). To achieve cascade enzymatic reactions, multiple chips can be stacked up with tube-less connections between inlets and outlets (**Fig. 3c**, Supplementary **Fig. S3**). 3D-printing offers obvious advantages over traditional chip fabrications, such as fast turn-around time, creation of 3D structures with complex architectures, and elimination of the chip bonding process.

**Figure 3.**
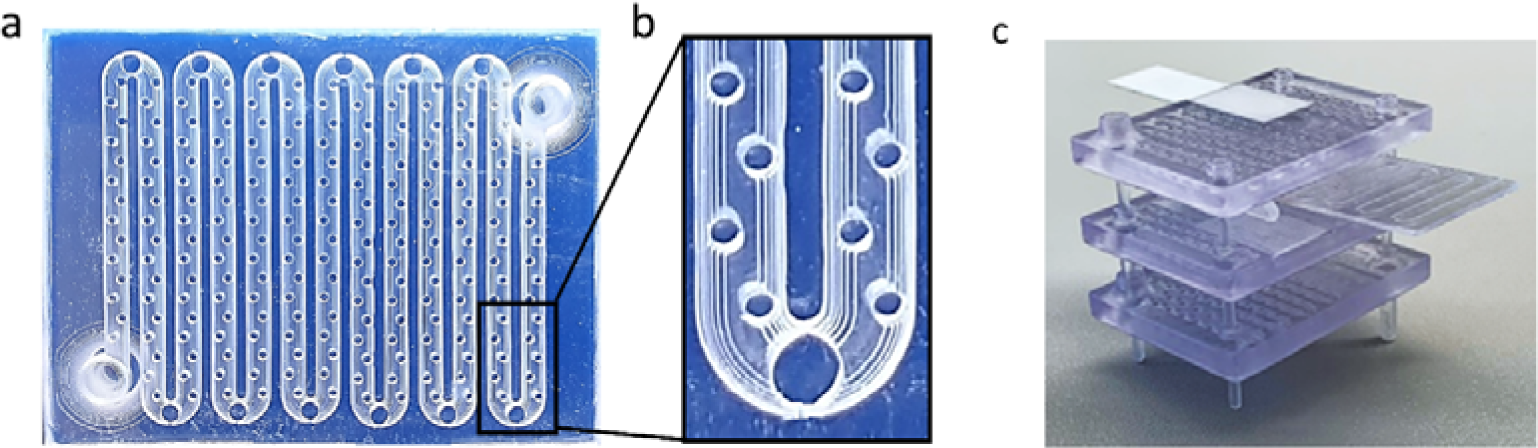
**a.** 3D-printed TurboMix chips. **b.** Enlarged image of the channel structure. **c.** Multiple chips stacked up with tube-less connection between inlet and outlets.

### Enzyme immobilization

Besides the chip configuration, the methods used to immobilize the enzymes to the chip surfaces are also critical for microfluidic bioreactors, as they significantly affect the amount, activity, and stability of the enzymes^22^. To explore the potential of enzyme immobilization on the printed resins, we initially investigated the utilization of chemicals to generate hydroxyl groups on the surface of the hydrophilic biocompatible resin. We tried various surface activation methods like combinations of oxidizing agents with sulfuric acid, but these methods showed detrimental impacts on the printed resin surfaces. They resulted in either resin dissolution or the formation of holes. Eventually, we devised a chemical approach to activate the resin surface using a combination of ammonium hydroxide and hydrogen peroxide, which enhanced the creation of active hydroxyl groups. Afterwards, we immobilized β-galactosidase enzyme on the chips by using the chemical linkers APTS and glutaraldehyde. Investigation of the X-gal intensity showed that the bioactivity of the immobilized enzyme was well maintained (Supplementary **Fig. S5)**. The Scanning Electron Microscopy (SEM) image of the channels revealed that this chemical treatment method did not compromise the integrity of the printed resin structure (**Fig. 4**).

**Figure 4.**
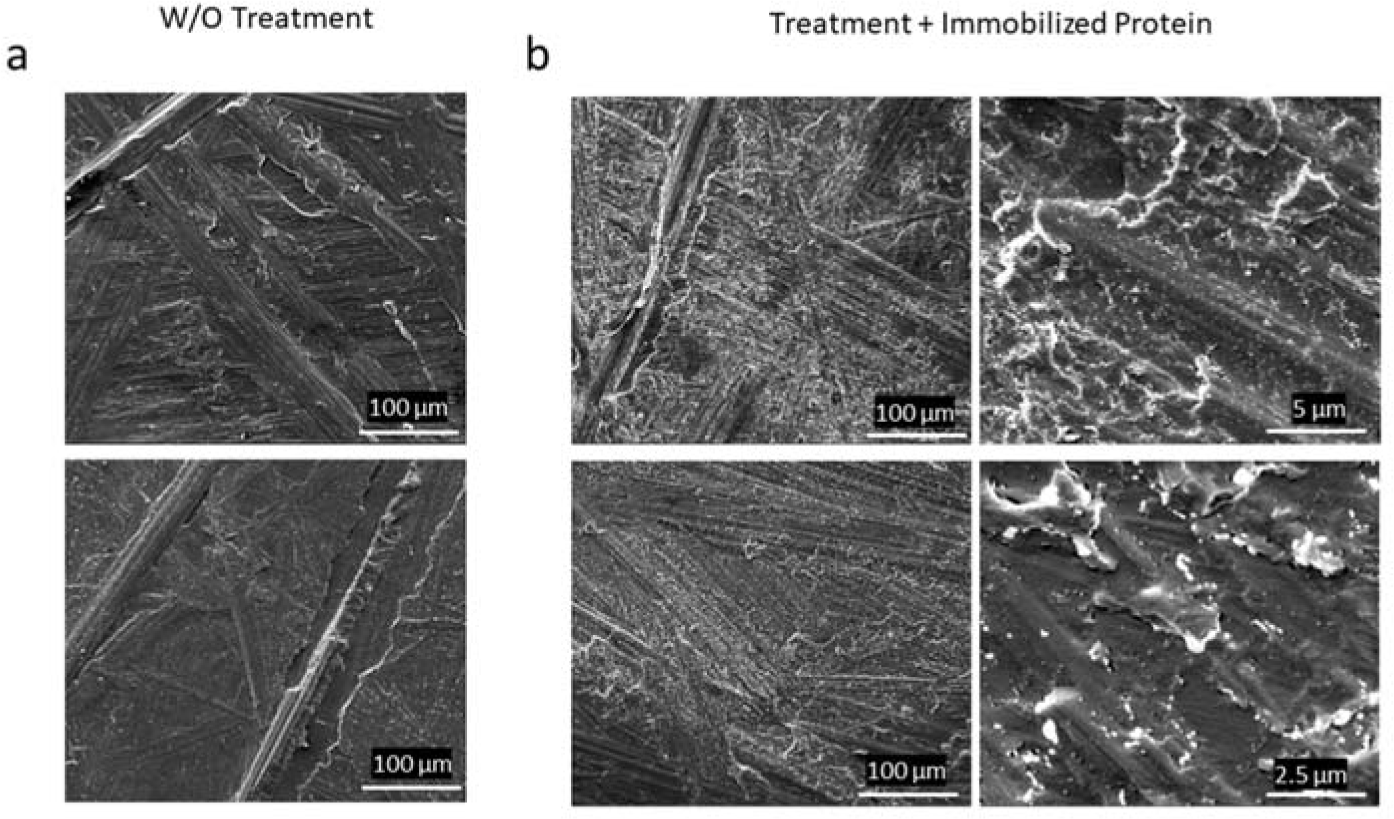
SEM images of the channel surfaces. **a.** Channel surface in its original state. **b.** The channel surface following activation and protein immobilization.

To further augment the enzyme concentration at the channel surfaces, we explored the possibility of generating supplementary branches on each active chemical group. With this aim in mind, we introduced PAMAM dendrimer nanoparticles to branch out from each stem of APTS molecules (**Fig. 5**). Dendrimers are three-dimensional branched tree-like structures, consisting of an ethylenediamine core, a repetitive branching amidoamine internal structure and a large amount of primary amines at terminals^23^. The utilized G5 PAMAM dendrimer particle has a diameter of 5.4 nm and contains 128 primary amine groups. These amine functional groups are easily accessible, ensuring highly efficient enzyme conjugation. Consequently, this strategy allowed for the potential of 128 active amine groups attached to each APTS molecule for enzyme immobilization, leading to a much improved enzyme concentration.

**Fig 5.**
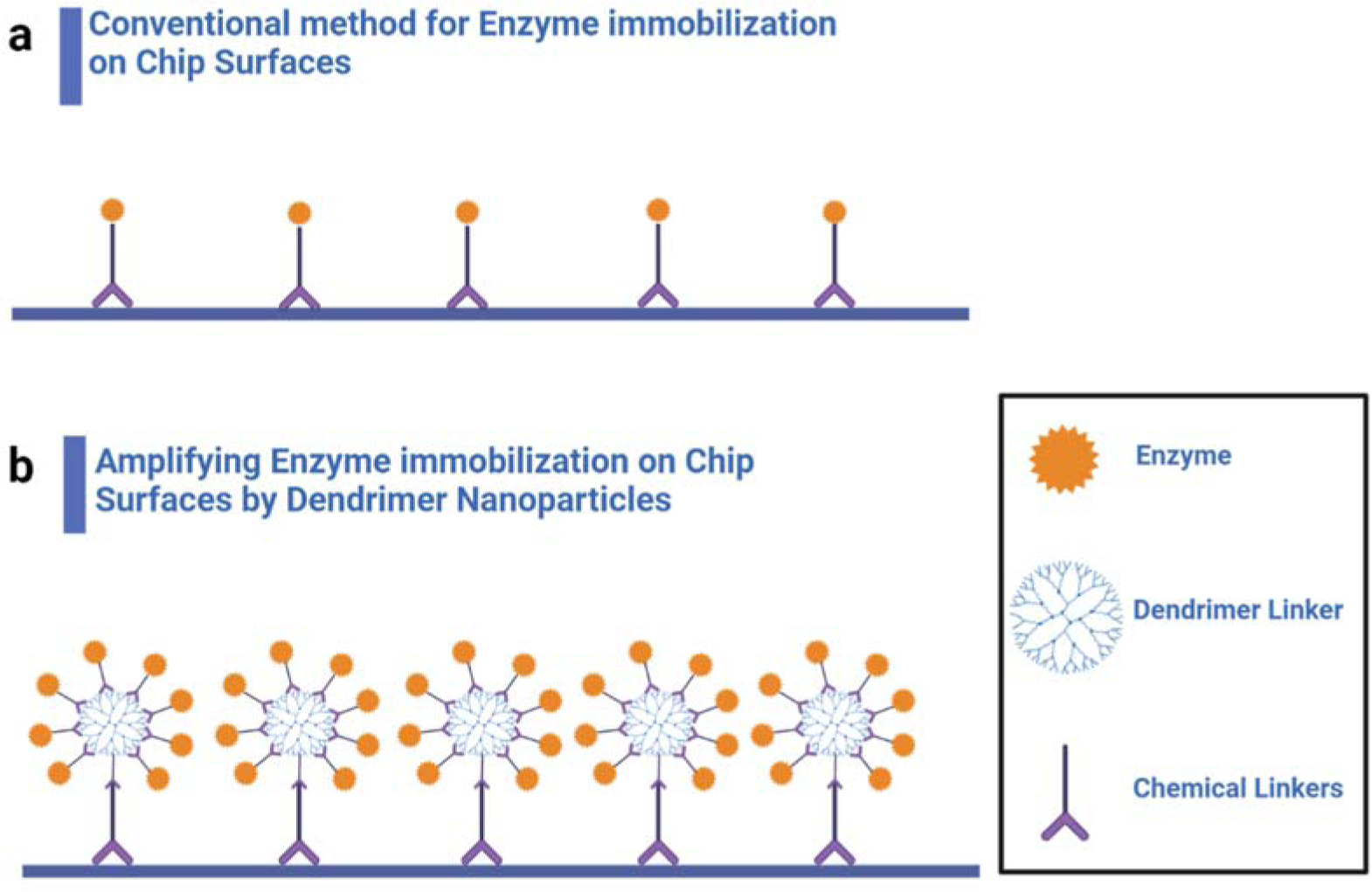
Enhanced method for protein immobilization on the 3D-printing resin surface. **a.** The conventional method is depicted, wherein only a single site is available for attaching proteins to each chemical linker on the resin surface. **b.** Incorporation of PAMAM dendrimer nanoparticles, which enables the generation of multiple branches on each attached chemical linker, thus offering increased protein attachment sites on the surface.

In light of these developments, we immobilized the GFP protein and β-galactosidase enzyme separately on the chips to assess the differences between the unmodified and modified approaches utilizing dendrimer nanoparticles. The results revealed a higher intensity of green fluorescent emission when dendrimers were employed (**Fig. 6a**), and the intensity of X-gal converted by β-galactosidase exhibited a five-fold increase with the presence of 25% PAMAM dendrimer (**Fig. 6b**). These studies showed that our chemical protocols in combination with the dendrimer coating presented a mild surface modification strategy and guaranteed the high loading of enzymes onto the microfluidic chip.

**Figure 6.**
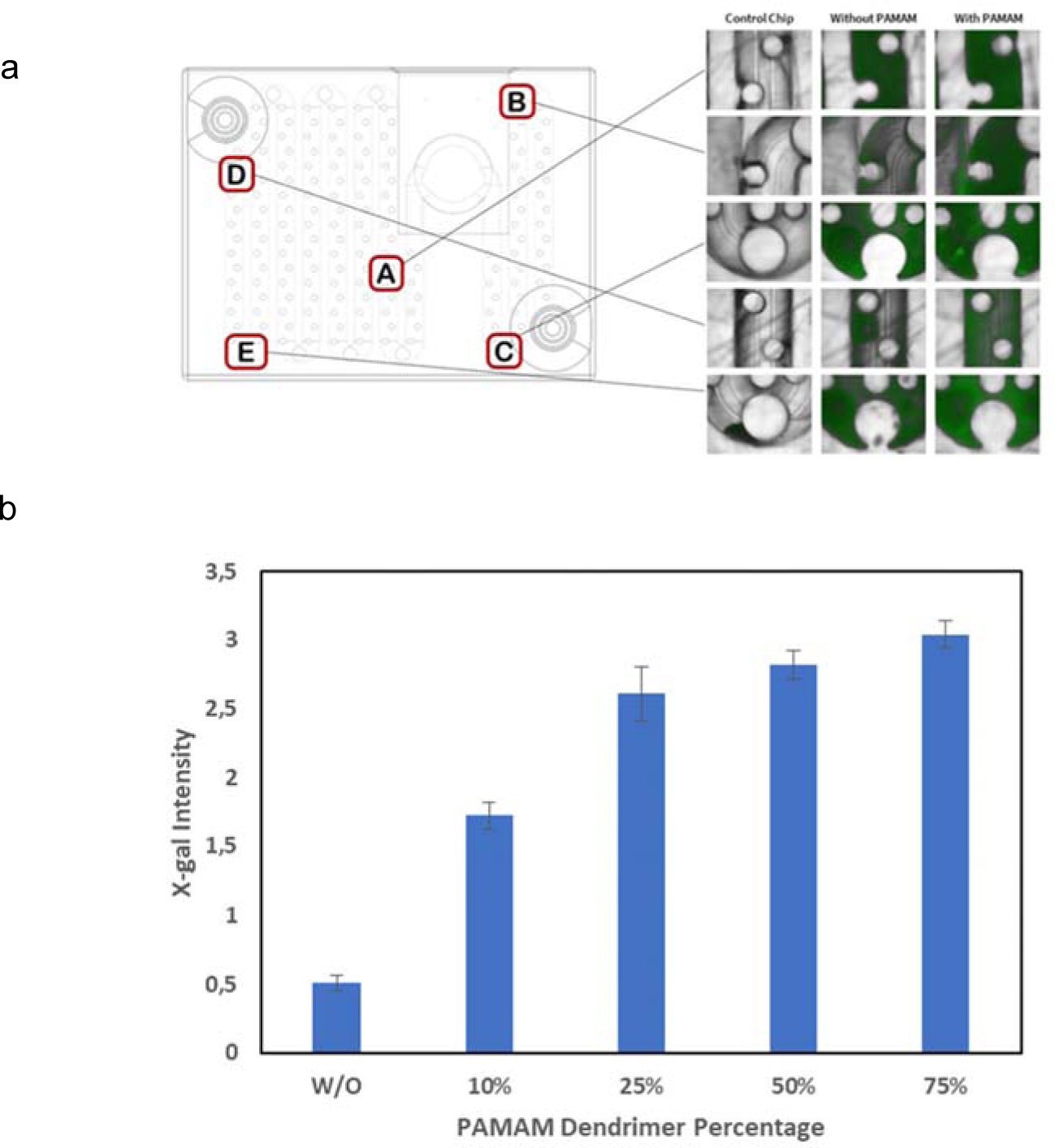
Effect of dendrimer coating on protein immobilization. **a.** Fluorescent microscopy images of the chips immobilized with GFP protein, revealing enhanced fluorescent emission in the chips with incorporated PAMAM dendrimers. **b.** β-galactosidase activity and X-gal intensity within the chips with different PAMAM dendrimers concentrations.

### Electron transfer to the enzymes

A lot of enzymatic reactions rely on cofactors such as NADH for electron transfer and hydrogen shuttle. These cofactors naturally exist in living cell factories and are oxidized during cellular respiration. However, in cell-free enzymatic reactions, the use of cofactors poses a significant challenge, including their short half-life in high temperatures or acidic/basic pH, plus their high cost^24,25^. Therefore, it is important to regenerate the cofactors from their oxidized form to make the cell-free metabolic systems more feasible for industrial use. Previously, several methods for cofactor regeneration have been tested, including photochemical, biological, and electrochemical processese^26,27^, whereas none of them can be readily adapted to microchips. To facilitate electron transfer in cell-free enzymatic reactions, hereby we innovatively introduced an electrode component into the microfluidic chip (**Fig. 7a**), and conducted investigations to explore two different approaches: direct electron transfer and co-factor (NAD(P)^+^) reduction (**Fig. 7b,c**).

**Figure 7.**
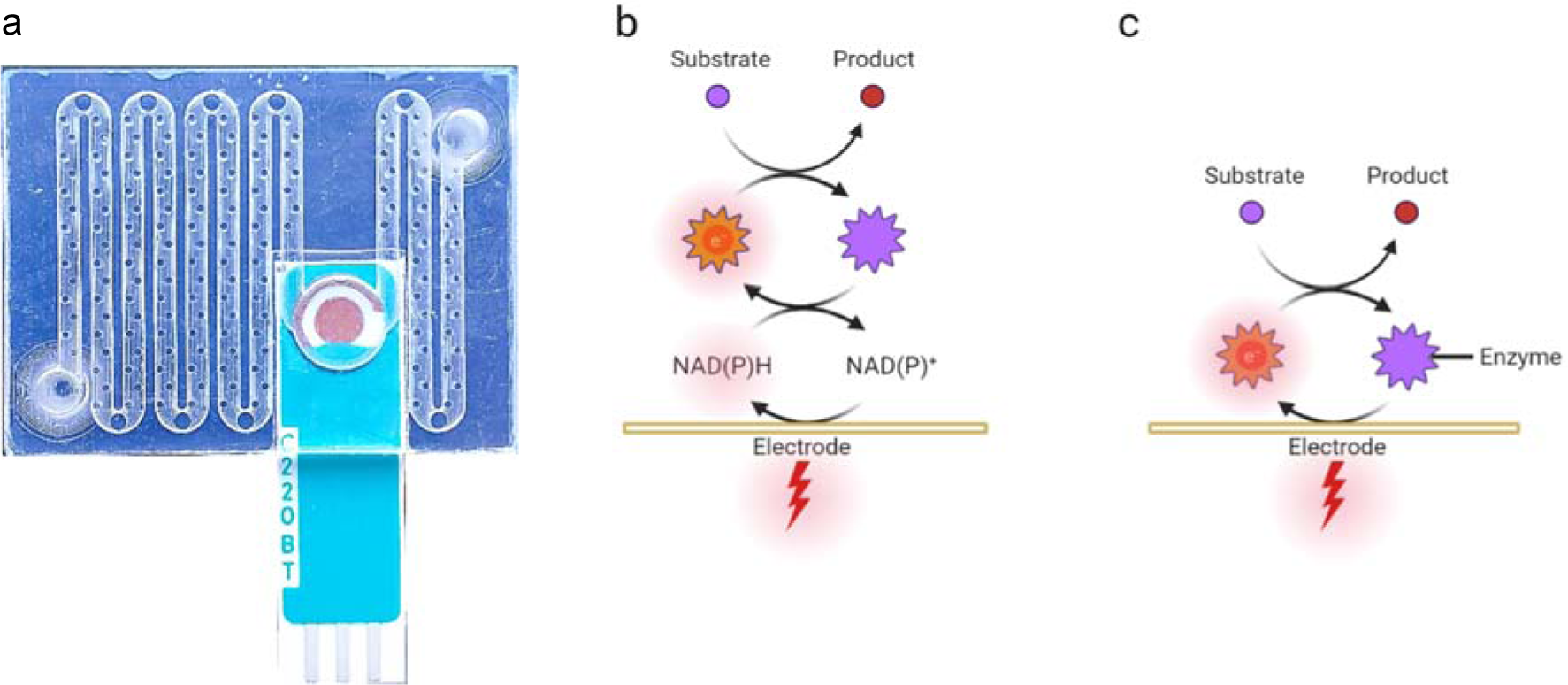
**a.** The electrode in chip setup. Electron transfer strategies to the enzymes. **b.** Indirect electron transfer: An electron-transferring cofactor, such as NAD(P)H, can undergo recycling from its oxidized form NAD(P)^+^ by an electrode. Subsequently, enzymes utilize the supplied electrons from cofactors to catalyze the conversion of substrates into products. **c.** Direct electron transfer: Electrons can be directly transferred to enzymes immobilized on the electrode’s surface, facilitating the conversion of substrates into products.

For the assessment of direct electron transfer, we immobilized LDH on the electrode surface and conducted reactions without NADH/NAD^+^ involvement. As shown in **Fig. 8a**, pyruvate was converted into lactate when negative voltage was applied, demonstrating the screen-printed electrodes were capable of directly transferring electrons to the enzyme. The amount of converted pyruvate was proportional to the negative potential. Moreover, we examined the extent of electrochemical transformation of pyruvate and lactate on the electrode in the absence of LDH enzyme, which shows that the concentration of pyruvate remained almost the same without LDH.

**Figure 8.**
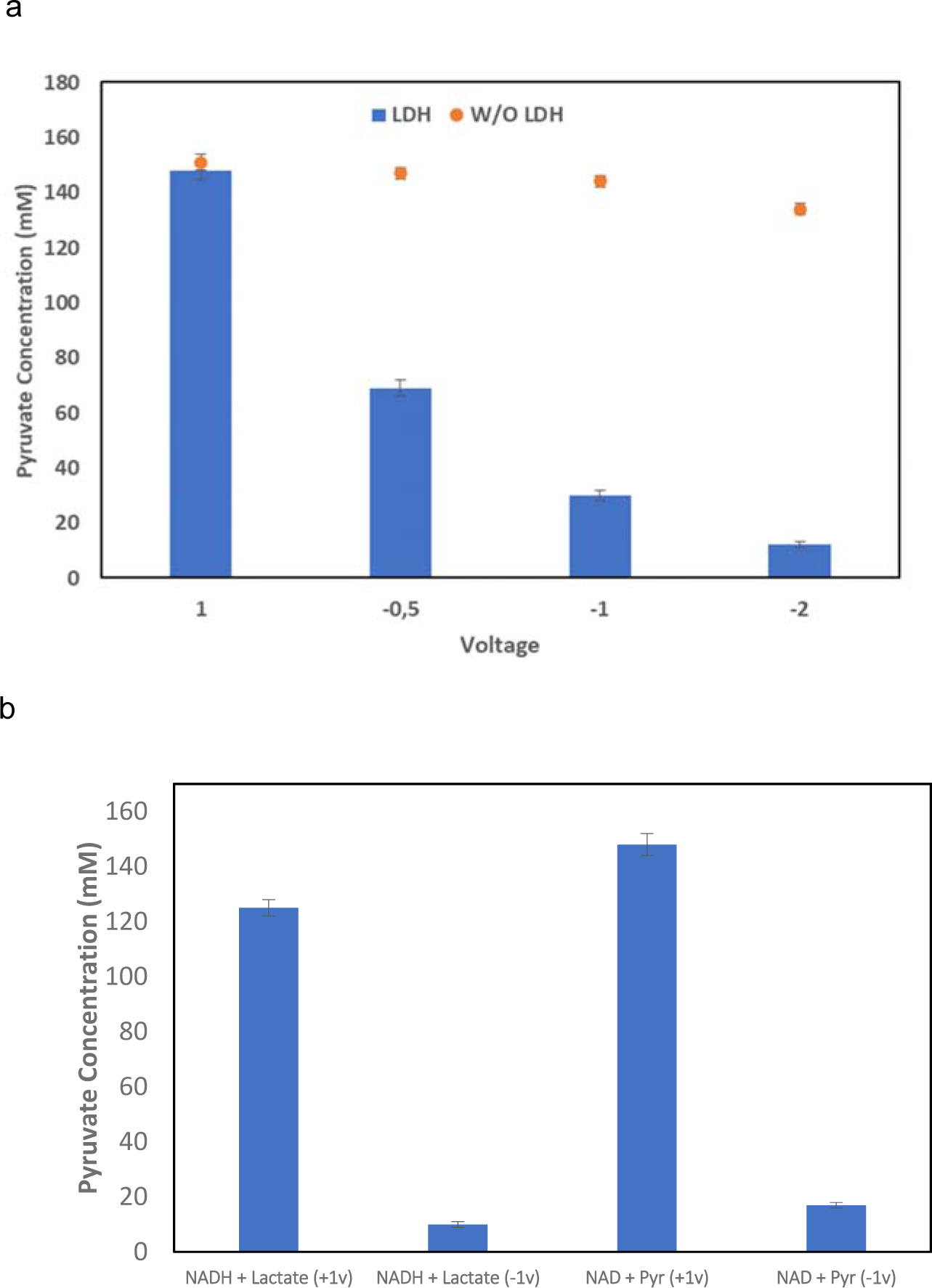
**a.** Direct electron transfer to LDH and pyruvate conversion to lactate by LDH in different voltages. **b.** Electron transfer to NAD^+^/NADH. Lactate and pyruvate conversion by LDH in different conditions. The impact of negative and positive voltages on the reaction in the presence of NAD^+^ or NADH was investigated. The conditions without LDH were studied as a control.

To regenerate NADH, we immobilized lactate dehydrogenase enzyme within 3D-printed chips and employed an active electrode component along with NAD^+^ or NADH. Subsequently, we evaluated various voltages and conditions while measuring the conversion of pyruvate. Notably, we achieved control over the reaction direction by applying positive or negative voltages, enabling the conversion of pyruvate to lactate or vice versa. We investigated the transformation of pyruvate into lactate using both +1 volt and −1 volt with 150 mM pyruvate and 15 mM NAD^+^ cofactor added to the reaction buffer. The findings demonstrated that at −1 volt, NADH was regenerated, and pyruvate was converted into lactate. We also explored the reverse reaction, converting lactate into pyruvate, by adding 15 mM NADH and 150 mM lactate to the solution and assessing pyruvate production at both +1 volt and −1 volt. Our results indicated that at +1 volt, NAD^+^ could be generated, and lactate was converted into pyruvate (**Fig. 8b**).

Furthermore, we fine-tuned the voltage, buffer composition, and minimal medium to ensure efficient electron transfer without causing any detrimental effects to the electrode. We observed that PBS (1x) was unsuitable for efficient electron transfer within the designed system, and its use significantly reduces the electrode’s half-life. We proceeded with diluted PBS (0.1x) and 20 mM tris-HCl buffers. Encouragingly, our results demonstrated that the electrode exhibited satisfactory performance with these buffers, even at higher voltages (±7 V), for a duration exceeding 24 hours. Notably, no detrimental impact on the electrode was observed over a span of 72 hours.

In short, both of these examined methods exhibited efficacy in facilitating electron transfer directly or indirectly by regenerating NAD(P)H within the system. Considering the enzyme immobilization capacity, we chose the co-factor NAD(P)^+^ reduction approach in the subsequent studies.

### Design of pathways for xylose consumption and CO_2_ fixation

To demonstrate that metabolic engineering can be implemented on the 3D printed microfluidic systems, in this study, we specifically focused on the conversion of lignocellulosic compounds that play a crucial role in second-generation fermentation. Xylose, an important five-carbon sugar derived from lignocellulosic materials, represents a promising renewable resource for biofuel and chemical synthesis, while *S. cerevisiae* is a preferred microorganism for ethanol and biofuel production^28,29^. Nonetheless, natural *S. cerevisiae* lacks the inherent capacity to ferment xylose and to assimilate inorganic carbon into biomass. While extensive efforts have been dedicated to genetically modifying yeast to enable xylose consumption^28,30^, we aimed to construct a xylose consumption pathway on chip to convert xylose into consumable molecules for *S. cerevisiae* without resorting to genetic modifications. In addition, the assimilation of CO_2_ in yeast and other microorganisms emerges as a captivating objective in the field of metabolic engineering, aiming to establish sustainable fermentation processes^31–33^. Considering that yeast releases CO_2_ as a by-product during its fermentation, we would also build a CO_2_ fixation pathway to achieve the prospect of neutral CO_2_ ethanol production.

In order to facilitate the design of a short pathway, we employed Cytoscape StringApp software to analyze the closest pathway to xylose conversion in *E. coli* for potential CO_2_ assimilation. To accomplish this, we constructed separate networks for xylose conversion, glycolysis, pentose-phosphate, and all proteins related to CO_2_ emission/fixation from *E. coli* (Supplementary **Fig. S6 a-d**). Subsequently, we merged these networks to identify the enzyme most closely associated with the xylose conversion pathway and possessing CO_2_ fixation capability (Supplementary **Fig. S6e**). Notably, the Gnd enzyme emerged as the closest enzyme with the ability to fix CO_2_, facilitating the conversion of Ru5P to 6-PGA by incorporating a CO_2_ molecule. Moreover, the Gnd enzyme exhibits oxygen resistance, making it well-suited for CO_2_ fixation without the necessity of anaerobic conditions, unlike RuBisCo. The enzymatic activity of Gnd relies on NADPH as a crucial co-factor^34,35^. To optimize the utilization of co-factors in the reaction, we could employ the electrode system to regenerate NADPH from NADP^+^, thereby reducing the overall co-factor requirement. In this regard, we opted to immobilize both Gnd and Rpe enzymes within the same chip. This strategy allowed for the efficient conversion of Ru5P to 6-PGA while minimizing the reversal of the reaction catalyzed by Rpe, which converts Ru5p back to Xu5p. Building upon these findings, we incorporated the complete Gnd–Entner–Doudoroff (GED) pathway into the xylose conversion pathway to further enhance its performance and overall efficiency. In total, six enzymes were selected to accomplish the two pathways, including XylA, XylB, Rpe, Gnd, Edd, and Eda enzymes (Supplementary **Fig. S6f**). With the modular metabolic pathways, the xylose can be fermented, and the intermediates are transported to natural *S. cerevisiae* and subsequently transformed into the final products (**Fig. 9**).

**Figure 9.**
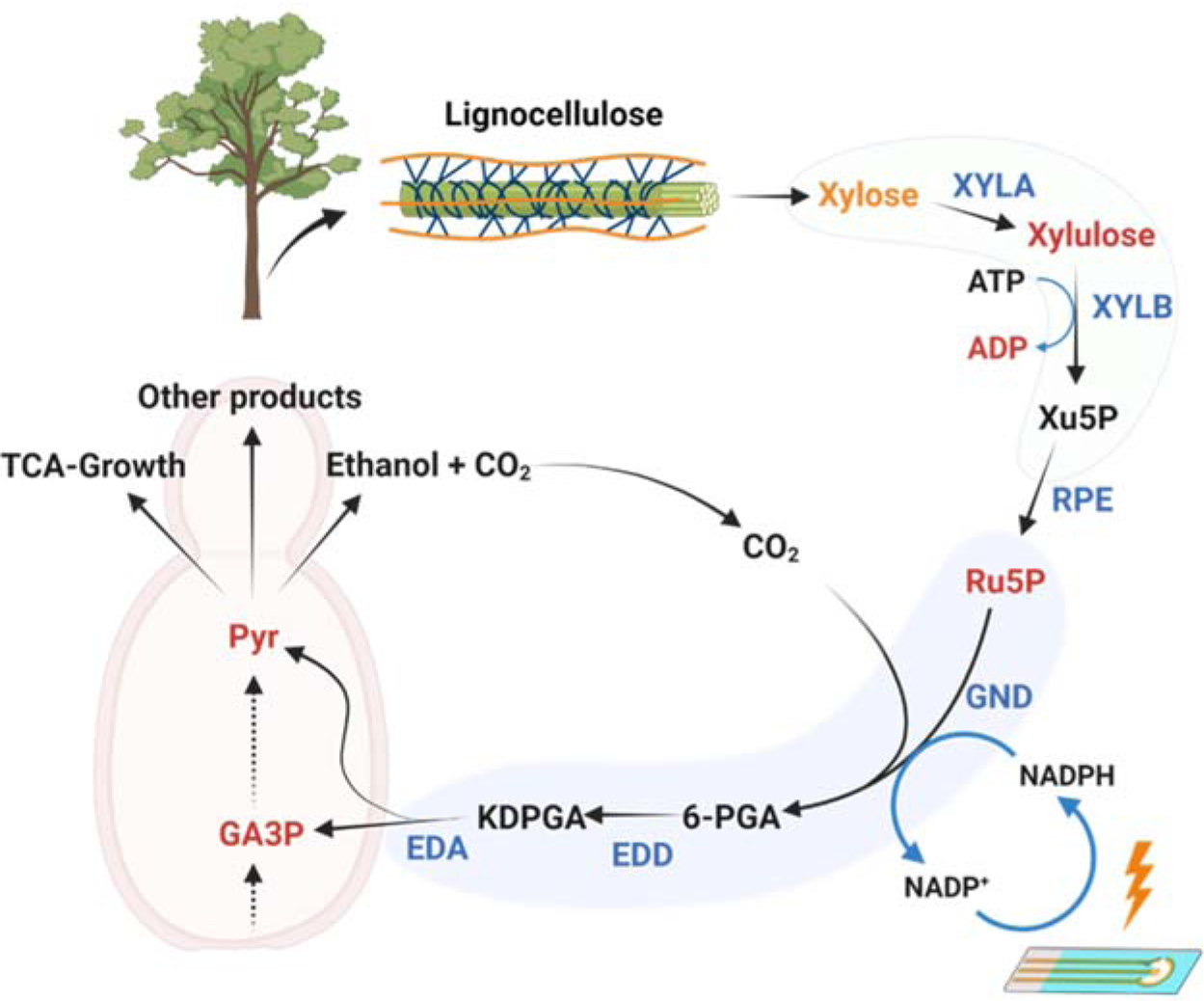
The selected pathways for xylose conversion and CO_2_ fixation. Illustration of the interface of the pathways and natural *S. cerevisiae*. Initially, xylose is consumed and converted into xylulose, xylulose 5-phosphate (Xu5P), and ribulose 5-phosphate (Ru5P) during the xylose conversion part. Subsequently, the Gnd enzyme catalyzes the conversion of Ru5P into 6-phosphogluconate (6-PGA), with the incorporation of a CO_2_ molecule and NADPH oxidation. Following this, the Edd enzyme transforms 6-PGA into 2-keto-3-deoxy-6-phosphonatogluconate (KDPGA). In the next step, the Eda enzyme is responsible for the production of pyruvate (pyr) and glyceraldehyde 3-phosphate (GA3P) from KDPGA. Notably, in the system, NADP^+^ can be reduced, enabling the recycling of NADPH for the Gnd enzyme by the electrode. The metabolites in red could be consumed by yeast.

### Utilized pathway and yeast culture

Subsequently, we proceeded to express and purify the enzymes encoded by the *xylA*, *xylB*, *rpe*, *gnd*, *edd*, and *eda* genes sourced from *E. coli*. These enzymes can play essential roles in the conversion of xylose to Ru5P, followed by the assimilation of a CO_2_ molecule catalyzed by the Gnd enzyme, ultimately yielding pyruvate and GA3P as the final products. The successful purification of the proteins was confirmed through SDS-PAGE (Supplementary **Fig. S7**), and subsequent enzymatic activity assays were conducted using conventional tube-based experiments.

We examined the yeast growth and uptake of metabolites using the minimal medium supplemented with different contributed metabolites (**Fig. 10a,b**). First, we tested the xylose conversion pathway, including xylulose, Xu5P, and Ru5P. Our findings indicated that yeast exhibited uptake and consumption of xylulose and Ru5P, while showing no growth on xylose and Xu5P. Considering the requirement for ATP during the conversion of xylulose to Xu5P in the xylose conversion pathway, we conducted yeast cultivation experiments using both ATP and ADP in the minimal medium. This investigation provided clarity on the growth behavior of yeast in the presence of these molecules. Remarkably, the results demonstrated that yeast exhibited no growth when ATP was provided as a carbon source. However, yeast demonstrated the capacity to consume ADP, which serves as a by-product of the pathway, thereby facilitating yeast growth. Afterwards, since the product of the Gnd enzyme was 6-PGA, we tested the yeast growth on this metabolite as a carbon source in the minimal medium. The results indicated that yeast could not grow on 6-PGA, potentially due to the yeast’s limited uptake capacity for this particular molecule. Consequently, we evaluated other metabolites within the GED pathway and cultured yeast on KDPGA and pyruvate+GA3P. Although yeast growth remained sluggish with KDPGA, it exhibited robust growth on Pyruvate+GA3P.

**Figure 10.**
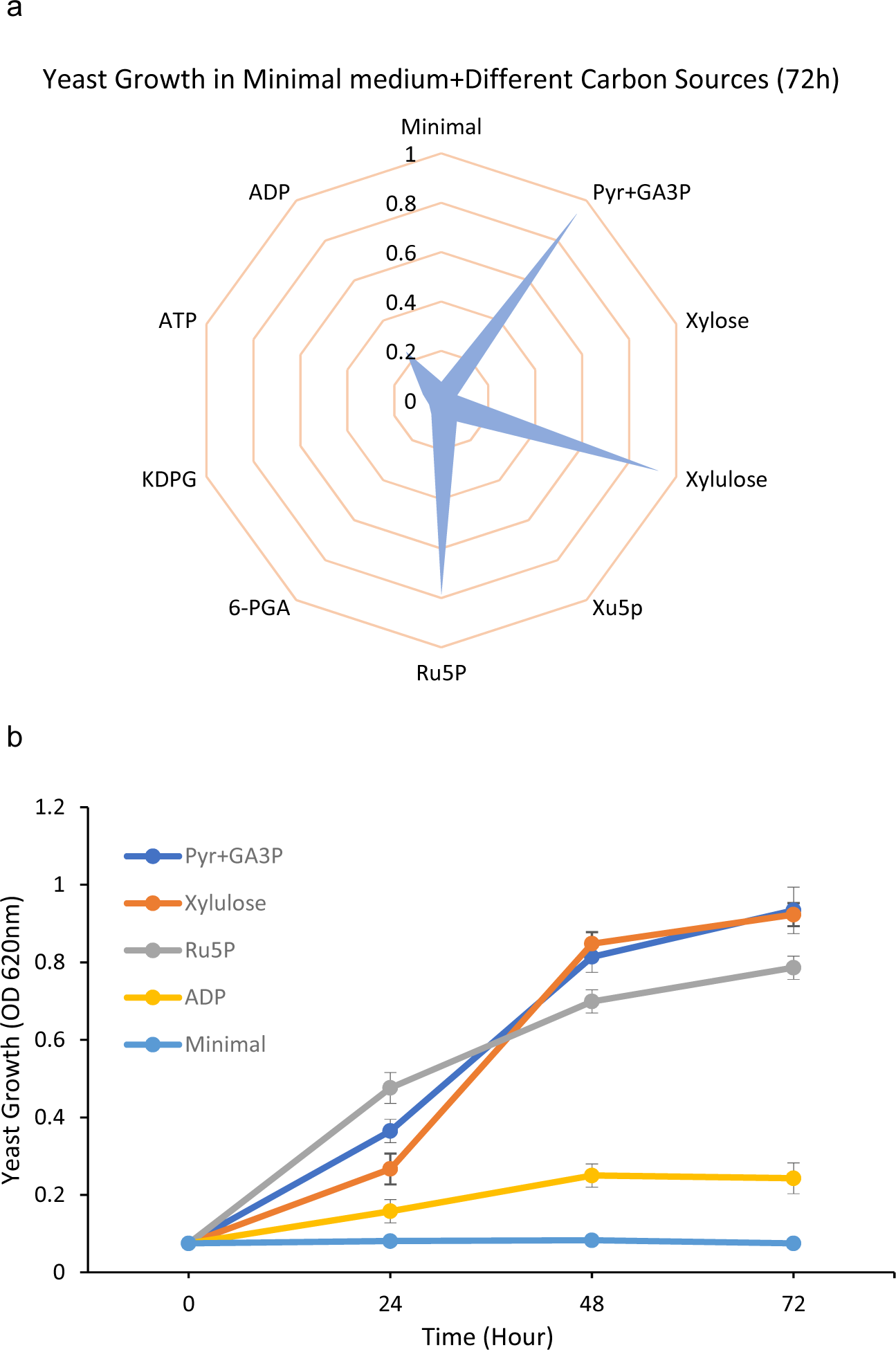
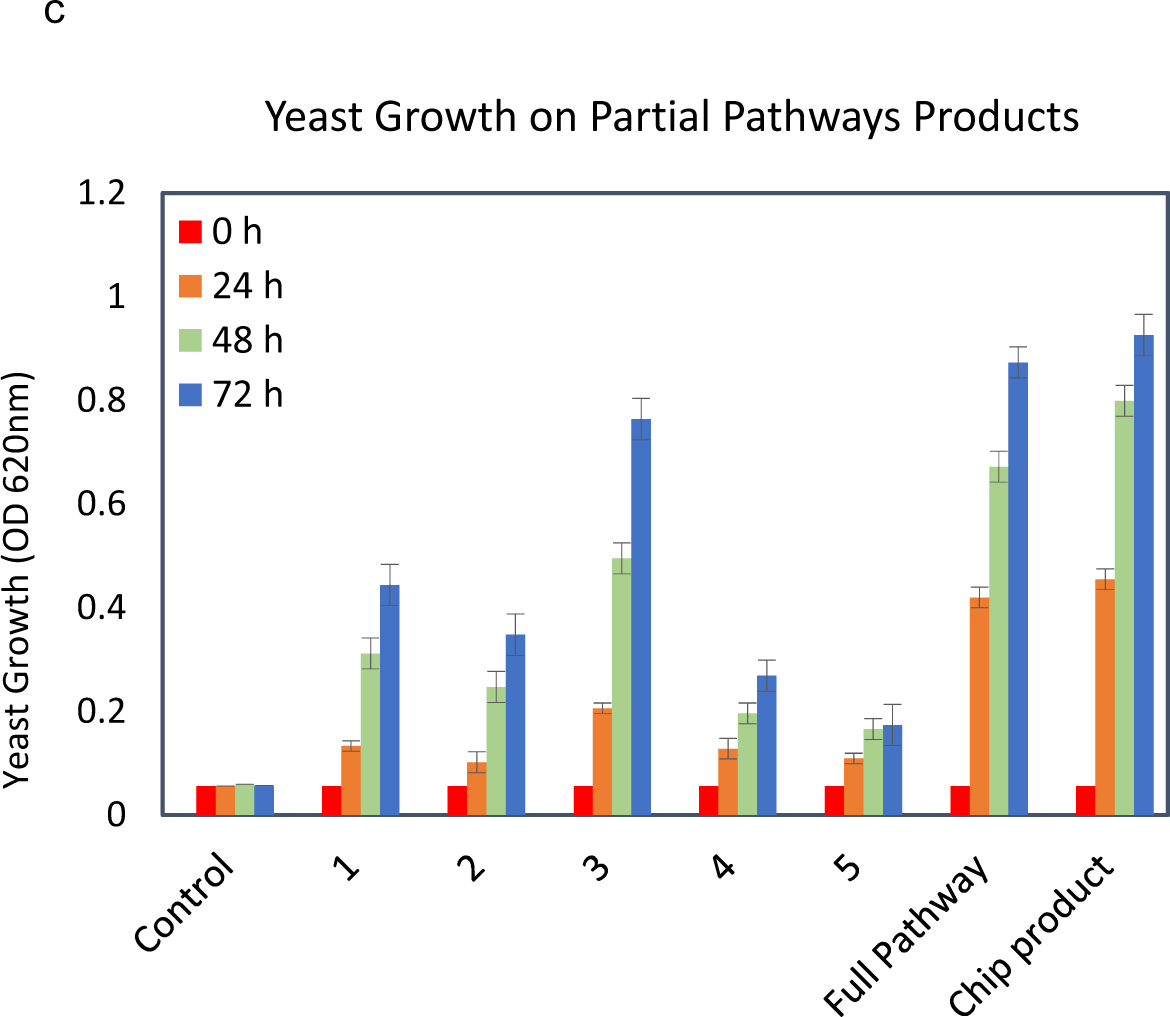
**a,b** Yeast growth was assessed in a minimal medium supplemented with different contributed metabolites from the employed pathway. The graph data presents the optical density (OD) readings at 620 nm, indicative of yeast growth levels. **c.** Yeast growth was evaluated based on the reaction products derived from various segments of the pathway and the complete pathway itself in tubes. The control condition involved a minimal medium devoid of any added carbon sources. The experimental groups were denoted as follows: Group 1 representing only XylA enzyme, Group 2 denoting XylA+XylB, Group 3 indicating XylA+XylB+Rpe, Group 4 comprising XylA+XylB+Rpe+Gnd, Group 5 featuring XylA+XylB+Rpe+Gnd+Edd, the full pathway encompassing all six enzymes in tube, and the complete pathway itself in microfluidic chips.

Furthermore, we assessed yeast growth by examining the reaction products generated from different segments of the pathway, as well as the entire pathway, in test tubes (**Fig. 10c**). The control condition consisted of a minimal medium without any additional carbon sources. In addition, we also utilized HPLC to measure the intermediates during the conversion of xylose (Supplementary **Fig. S8**). Our findings indicate that because the enzymes in the pathway are reversible, we couldn’t achieve a 100% product yield for each pathway segment. As the results show, there was a decrease in growth in the segments where the final enzymes XylB, Gnd, and Edd were involved. This decrease may be due to yeast not fully utilizing these metabolites, and the observed growth might be based on the consumption of intermediate metabolites that can be readily used. The enzymatic activity assays proved that the enzymes selected for the two pathways were fully functional.

### Integration of the 3D-printed microfluidic platform

In this investigation, we developed a 3D-printed microfluidic platform to execute the two cell-free metabolic pathways. The integrated system is shown in **Fig. 11**. Three reaction chips were stacked on top of each other to enable multi-step cascade enzymatic reactions, with two enzymes immobilized in each chip. We meticulously designed and incorporated specific inlets and outlets for the seamless interconnection of various chips, ensuring leak-free operation throughout the implementation without using tubes. Furthermore, a heating part was integrated at the base of the chips to ensure the requisite temperature for the desired reactions. Because of the thin bottom of the chip (1 mm), reaching the desired temperature took place in less than 1 min. As for the electrode component, we developed a chamber that featured a gap for electrode insertion, accompanied by a small chamber positioned atop the electrode. Moreover, as an innovative concept, we contemplated employing the 3D printer to fabricate a yeast growth chamber, which could be linked to the system. Additionally, in order to enhance CO_2_ gas concentration into the buffer, we incorporated a gas chamber within the system, thereby enhancing the capture of CO_2_ emitted from yeast fermentation within the yeast growth chamber or added CO_2_ gas into the system for subsequent assimilation. As such, we managed to make a self-sustaining, continuously operating system for on-chip metabolic engineering. In addition to cell-free metabolite production, we have demonstrated the capability to establish connections between the chip and viable microorganisms and diverse cell types. This novel approach enables the production of consumable or intermediate metabolites for these cells, as well as the conversion of their products into other molecules.

**Figure 11.**
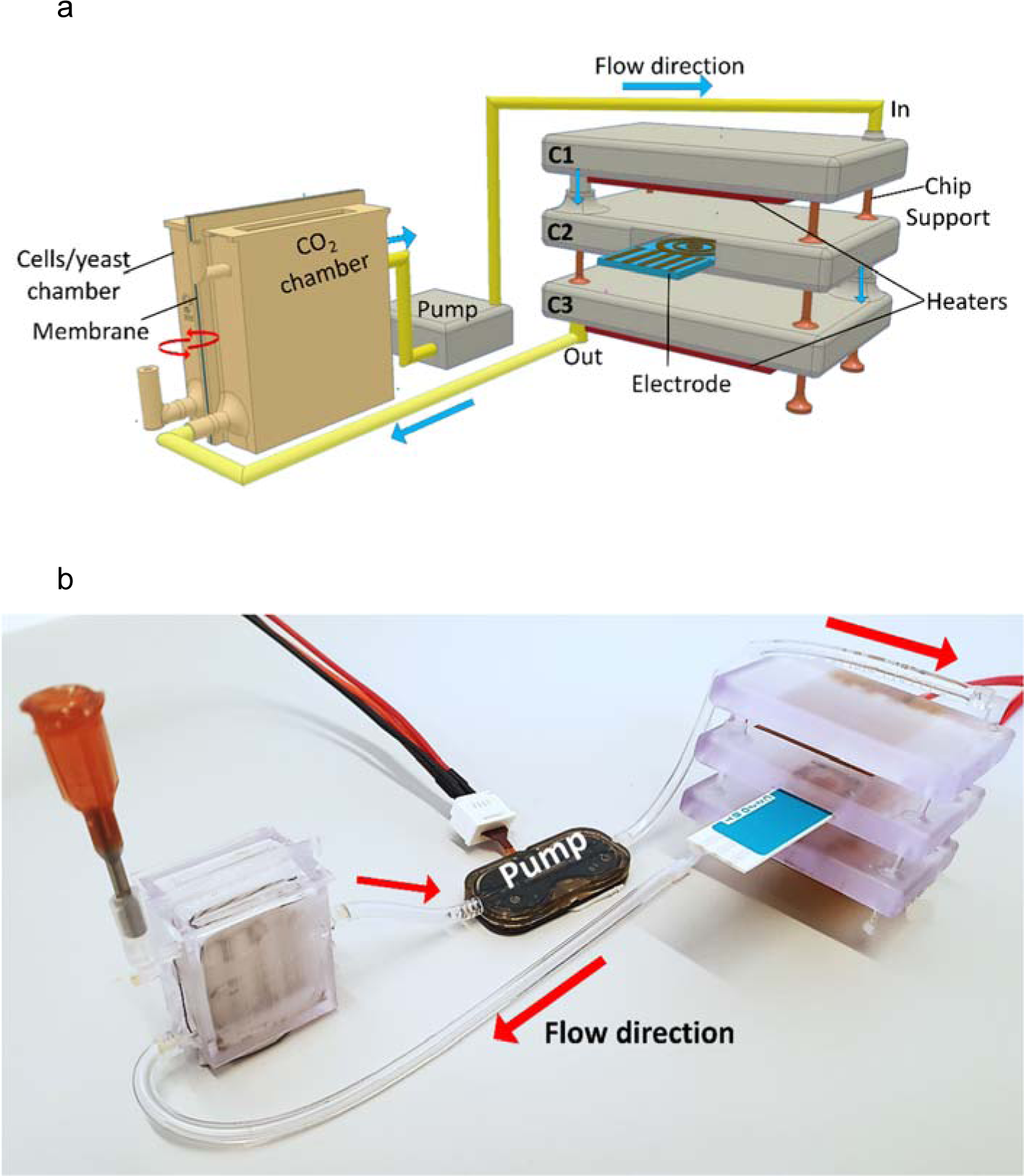
**a.** Schematic and **b.** real image of the integrated 3D-printed microfluidic platform.

### Yeast metabolic engineering on the chip

Given the yeast’s capability to consume xylulose and Ru5P as intermediate metabolites, we implemented two distinct strategies to link the microfluidic chips with the yeast growth chamber: a continuous system and a batch-to-batch xylose conversion system. In the continuous system, we established the connection between the chips and the cell chamber, inoculated the yeast, and filled the system with the minimal medium for the enzymatic reaction. In this setup, xylose was successfully converted into subsequent metabolites, enabling yeast to grow on intermediate metabolites prior to pyruvate and GA3P production. However, due to the comparatively slower growth rate of yeast relative to the enzymatic reactions occurring within the chip system, complete consumption of the intermediate metabolites was not achieved. Nevertheless, the chip system effectively facilitated the conversion of these intermediates, resulting in the production of pyruvate+GA3P.

In the batch-to-batch system, we operated the chip system for a 24-hour period before opening the valve connecting it to the yeast chamber. This strategy allowed the yeast to consume the final metabolites of the designed pathway. Although there were no significant differences observed between the two designed systems within the 72-hour period, the batch-to-batch system exhibited faster yeast growth in 24-hour (**Fig. 12**). Ultimately, our conclusive findings demonstrate that the designed system enables yeast to grow on a xylose carbon source while virtually achieving a relatively CO_2_-neutral fermentation process, all without requiring any genome manipulation.

**Figure 12.**
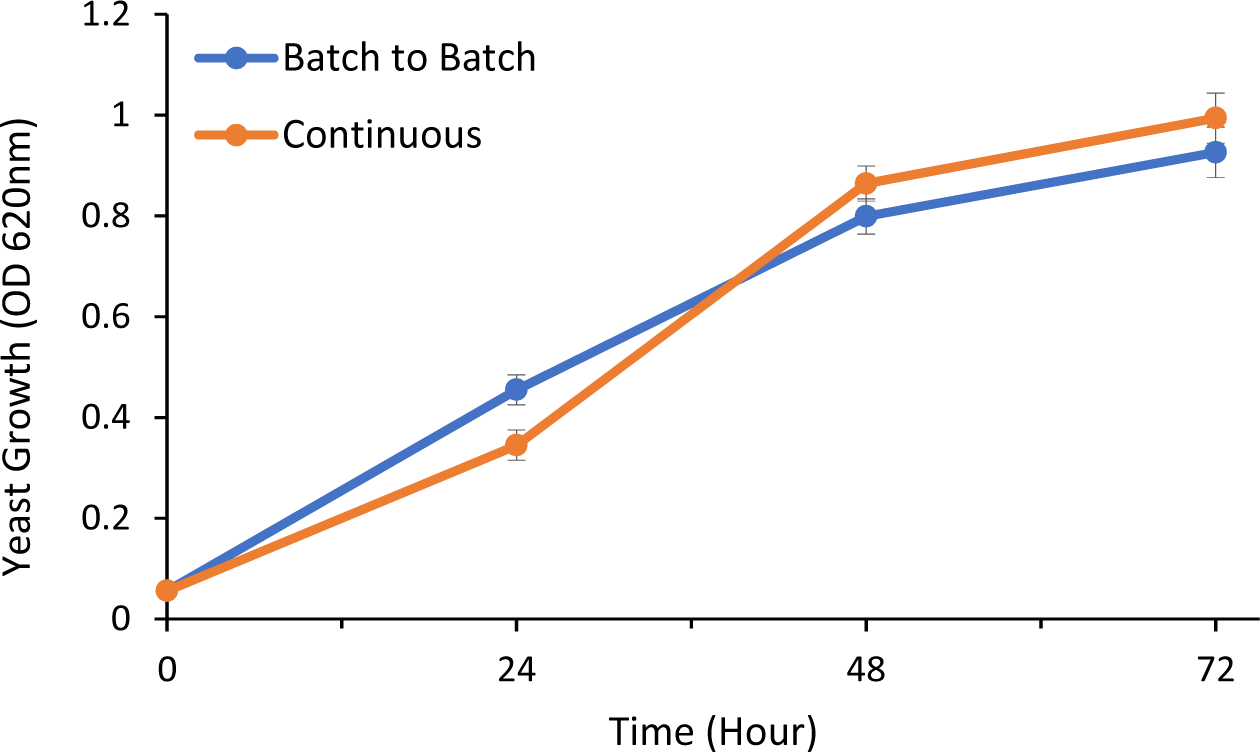
Yeast growth evaluation using the continuous system and the batch-to-batch system.

### Enzyme stability in the chip

To benchmark the stability of the enzymes within the microfluidic chips, we conducted a comparative analysis of the enzyme activity between the chips and conventional tubes. Specifically, we examined the activity of the designed pathway over a period of 7 days, both within the chips and the tubes. In the chip experiments, we maintained a continuous flow, changing the reaction buffer every 24 hours. For the tube experiments, the buffer was replaced with a fresh one using an Amicon column filter with a 3 kDa cut-off, and the reaction was measured using the same methodology.

The yeast growth findings demonstrated that the enzymes within the chips retained their activity for over 7 days, highlighting their remarkable stability (**Fig. 13**). In contrast, the enzyme activity in the tubes diminished within 48 hours. This comparison underscores the superior endurance of enzyme activity within the microfluidic chips, highlighting their potential for long-term enzyme stability and functionality.

**Figure 13.**
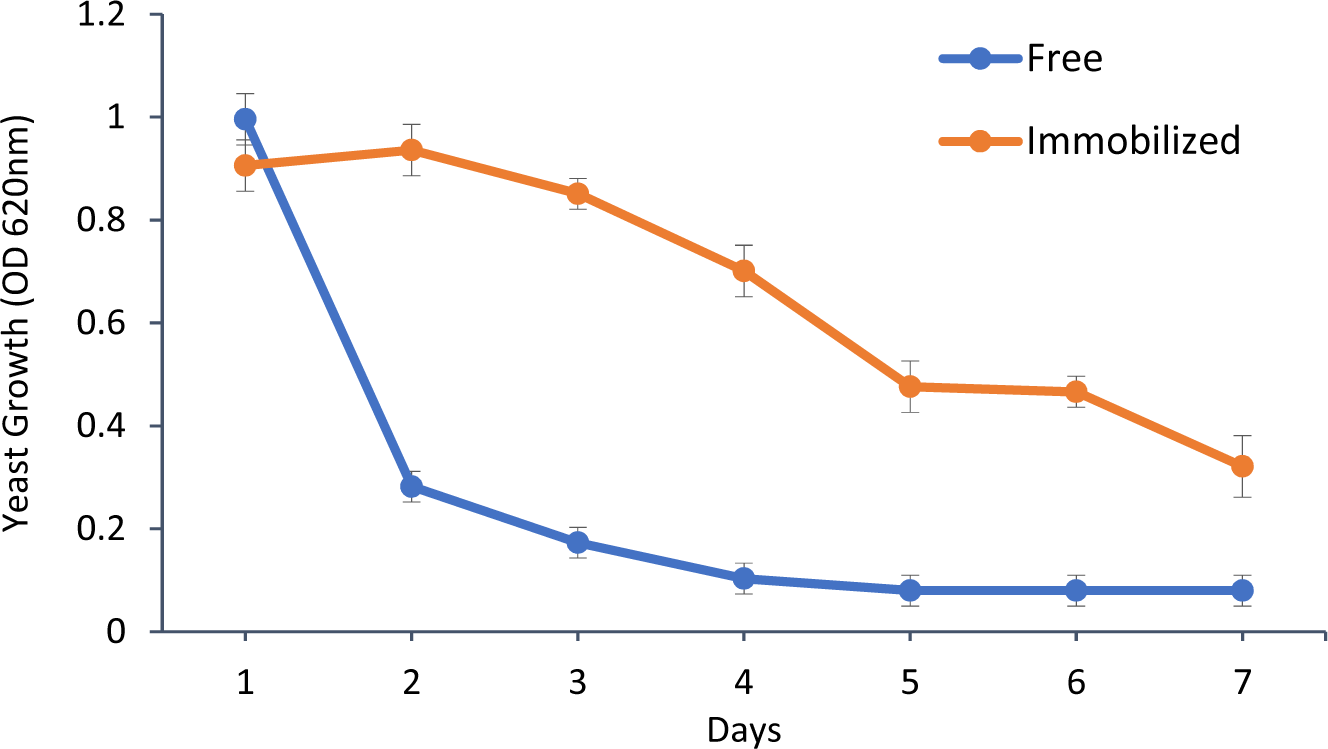
Yeast growth monitoring to assess the enzyme stability in free and immobilized forms.

## Conclusion

The development of new strategies for efficient metabolic engineering is essential to overcome challenges associated with conventional genome editing approaches^5,8,36^. This study addressed these limitations by utilizing a 3D-printed microfluidic platform for metabolic engineering. To achieve efficient mixing for enzymatic reactions, we designed a snake-like channel with irregular barrel configuration, and fabricated the chip by employing high-resolution 3D printing. We also managed to chemically activate the printed resin surface and immobilize enzymes in the channels. We showed that the amount of immobilized enzymes was significantly enhanced by coating dendrimer nanoparticles on the surface. Moreover, we established a new co-factor regeneration approach by inserting a gold screen-printed electrode in the microfluidic chamber to facilitate electro transfer in redox reactions. The study then focused on the metabolic engineering of *S. cerevisiae* on the microfluidic chip platform. Xylose consumption and CO_2_ fixation pathways were implemented to enable yeast to utilize xylose as a carbon source and assimilate CO_2_, leading to a virtually carbon-neutral fermentation process. By utilizing lignocellulosic five-carbon sugars and redirecting the CO_2_ released during fermentation, the work demonstrated the potential for sustainable biofuel and chemical production by the designed system.

To the best of our knowledge, this is the first study with the combination of 3D-printing technology, microfluidic chip design, and metabolic engineering strategies. Our research presented a facile way to construct separate pathways in chips for efficient metabolite production in a fully cell-free system. In addition, the chip system can also be connected seamlessly to a cell-factory fermentation system to form a tandem conversion system. We envision that the 3D printed microfluidic platforms hold promise for advancing the field of biotechnology and enabling more efficient and environmentally friendly production processes.

## Acknowledgements

This work was supported by Novo Nordisk Foundation of Denmark, grant NNF21OC0066562. We thank Dr. Ehsan Ansari Dezfouli for help with the SEM images, and Dr. Mohsen Mohammadniaei for help with the chemical methods.

## Author contributions

Seyed Hossein Helalat: Conceptualization, Methodology, Investigation, Writing- Original draft, Writing – Review & Editing. Islam Seder: Methodology, Investigation, Writing- Original draft, Writing – Review & Editing. Rodrigo Coronel Téllez: Methodology, Investigation, Writing- Original draft, Writing – Review & Editing. Mahmood Amani: Methodology, Investigation, Yi Sun: Conceptualization, Supervision, Writing- Reviewing and Editing, Funding acquisition.

