## Supplemental information for "Metabolic Engineering on a 3D-Printed Microfluidic Platform: A New Approach for Modular Co-Metabolic pathways"

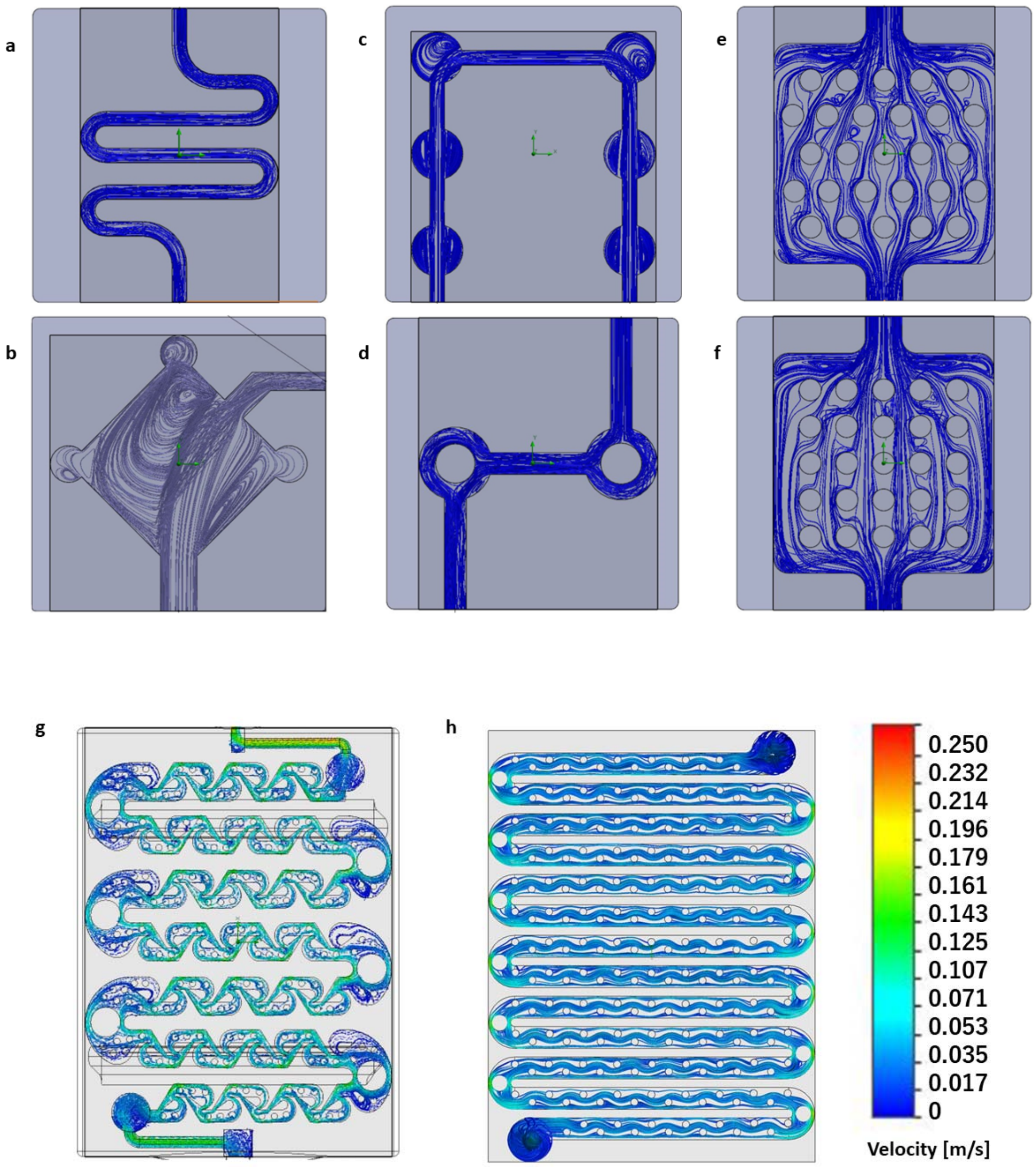


**Figure. S1 Various chip designs. a.** snake-like, **b.** circle-end diamond, **c.** bubbly, **d.** two-square, **e.** skewed-column, **f.** arranged-column, **g.** Tesla shapes, and **h.** TurboMix.

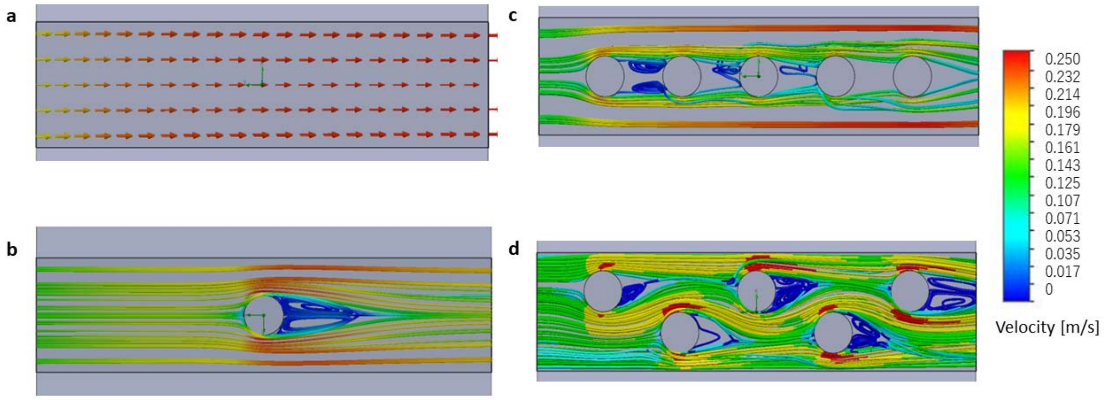


**Figure S2. Flow simulation of effect of barriers in microchannels. a.** Normal flow in a channel; **b.** Effect of a barrier on flow; **c.** Effect of tandem barriers; **d.** Effect of zigzag tandem barriers.

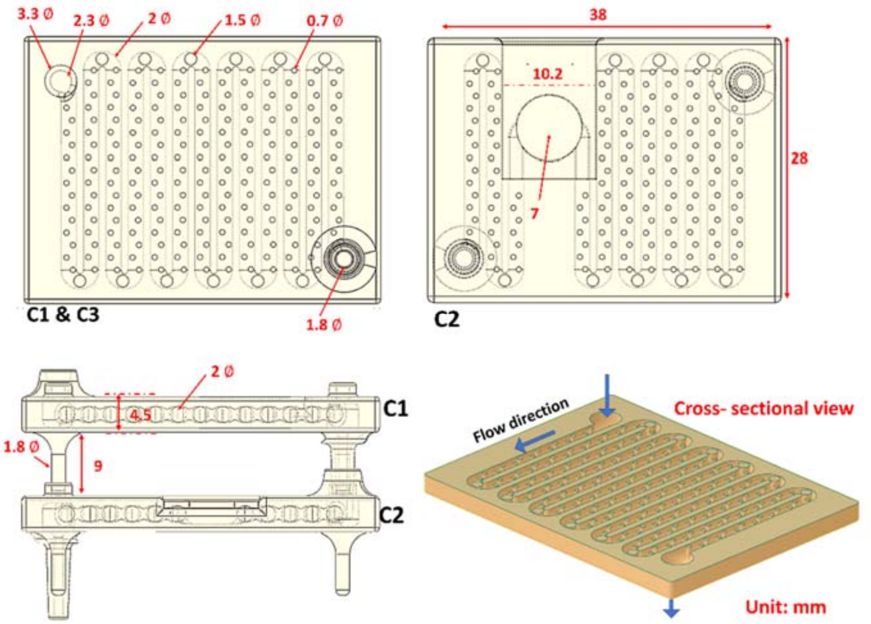


**Figure S3.** Detailed dimensions of the TurboMix chip.

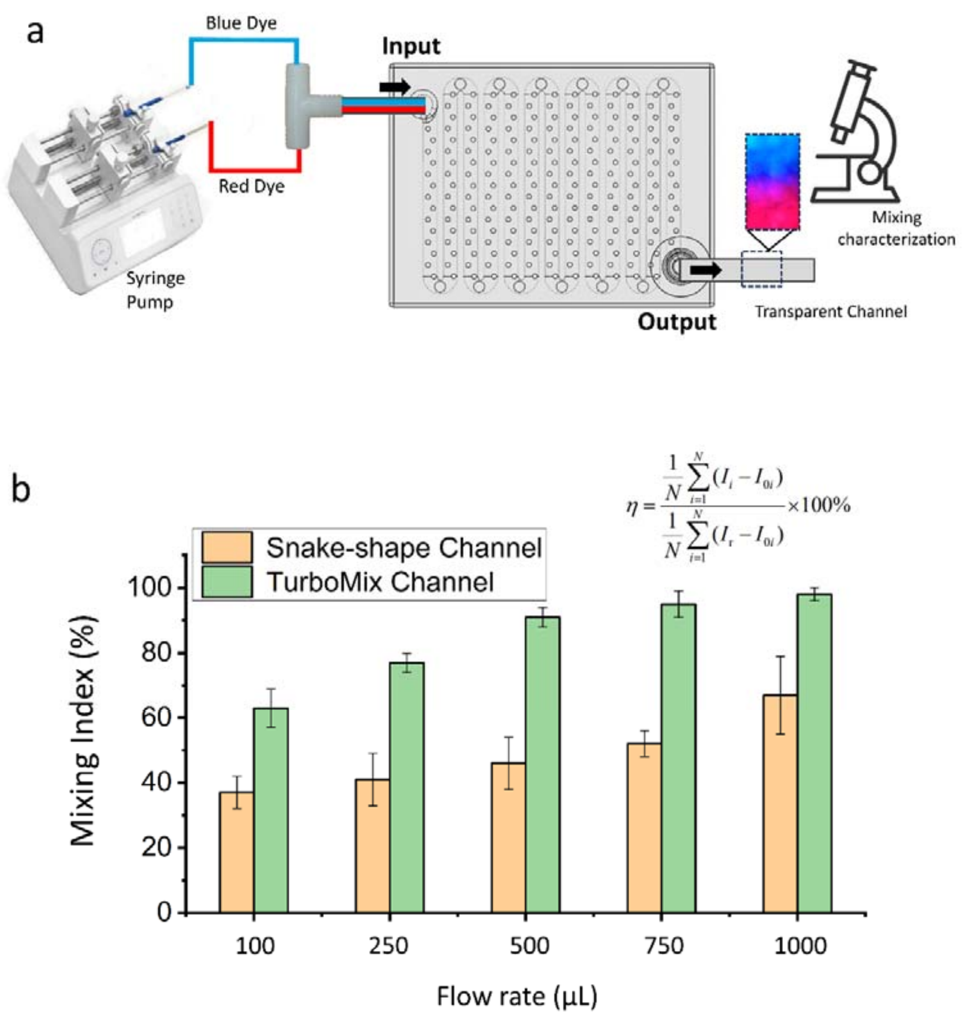


**Figure S4. Mixing characterization and reaction efficacy for the chip. a.** Experimental set-up. **b.** Mixing index. During the mixing, images were captured at a rate of 1 fps. Subsequently, the captured images were analyzed using the Octave software, and the mixing efficiency (η) was obtained using the mixing equation (right of fig.S4 b), where *I* is the normalized gray scale value, *I*i is the value of each pixel at the ith pixel of the captured image, *I*r is the normalized value of the reference image in the fully mixed state, N is the number of pixels, and *I*0i is the normalized value at the *i*th pixel in the initial state, i.e., without mixing. Thus, we obtain η = 0% for the pre-mixed condition and η = 100% for the completely mixed condition.

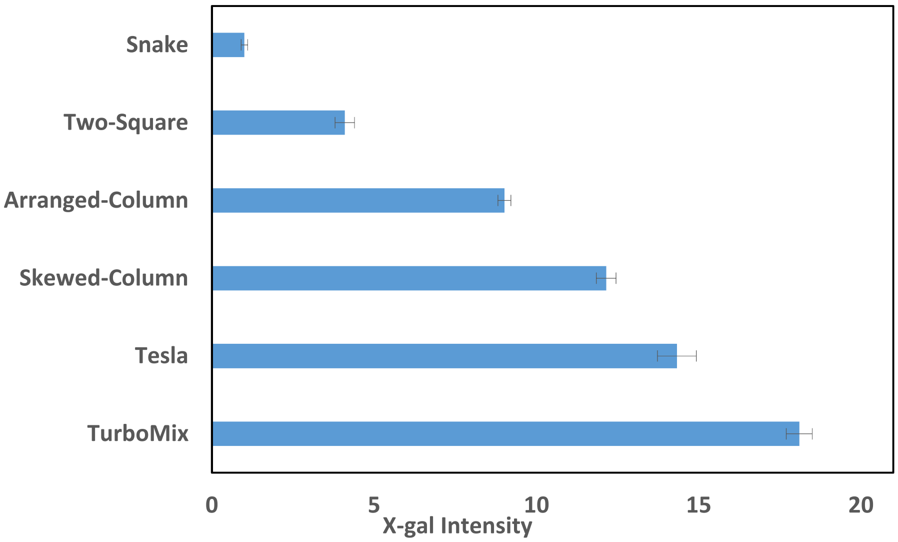


**Figure S5. β-galactosidase activity in channels with different shapes.** The enzyme β-galactosidase was immobilized on the microfluidic chips with various configurations. The color change of X-gal was observed in the channels for the β-galactosidase assay.

a. Glycolysis

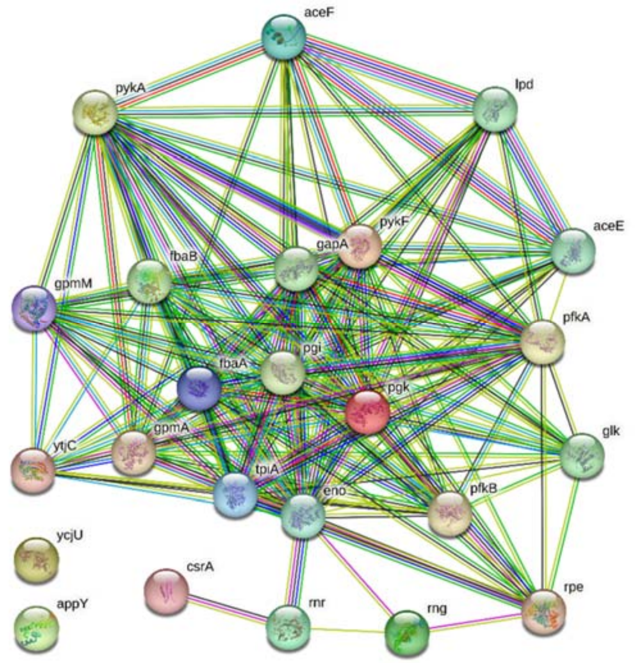


b. Pentose phosphate

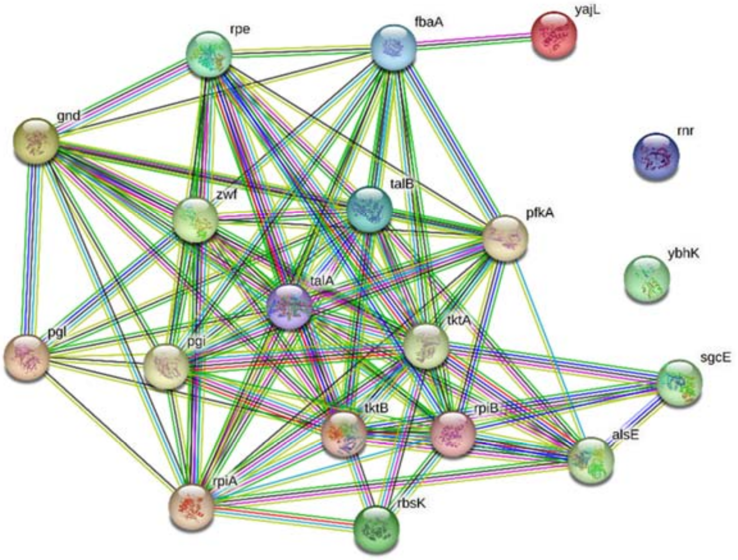

c. $CO_2$ emission/fixation

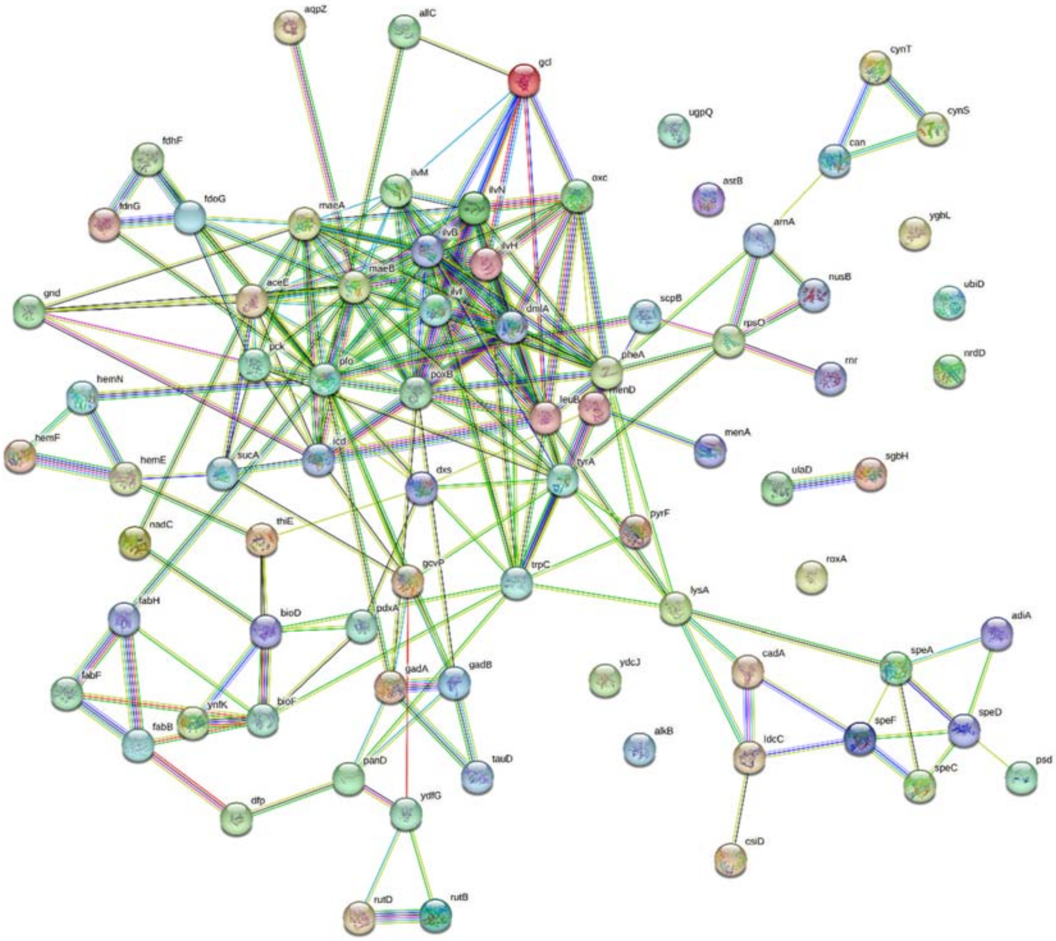

d. Xylose conversion

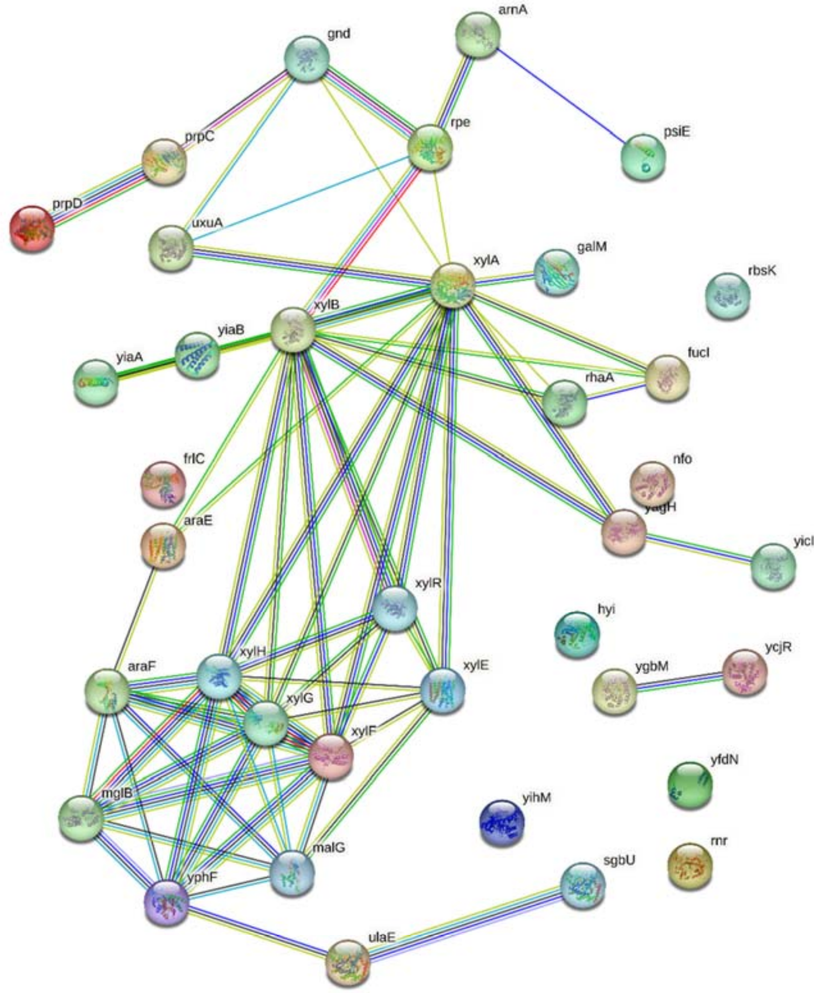

e. Merged

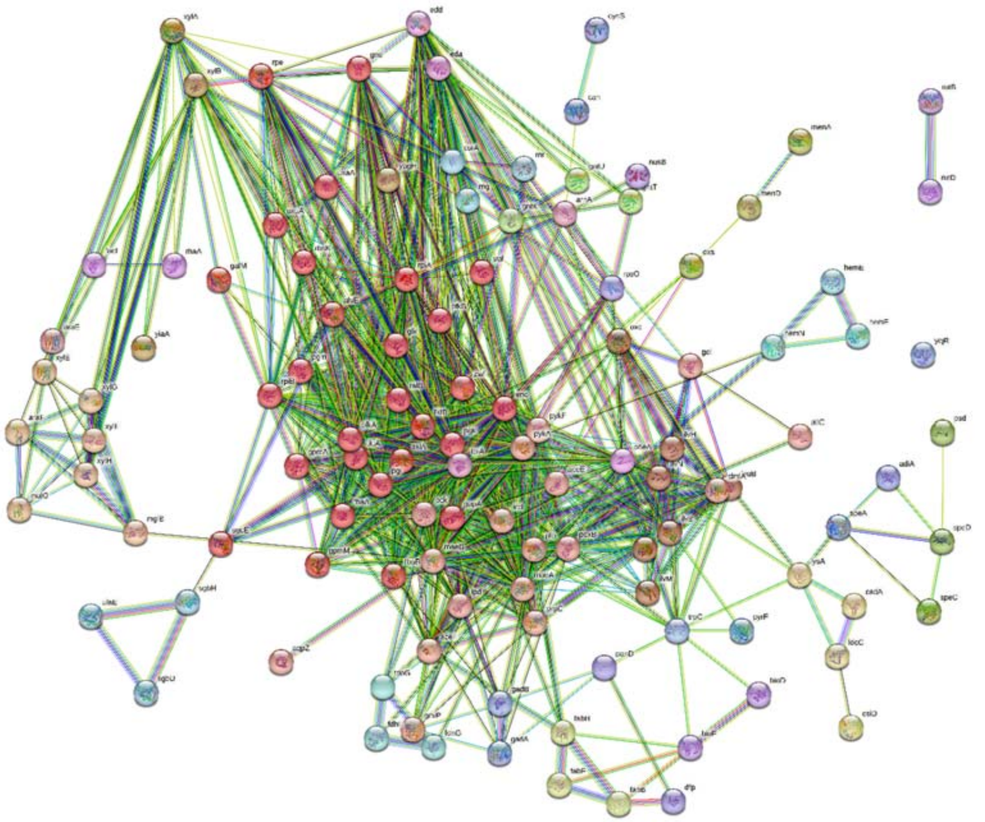

f. Selected pathways from the merged network

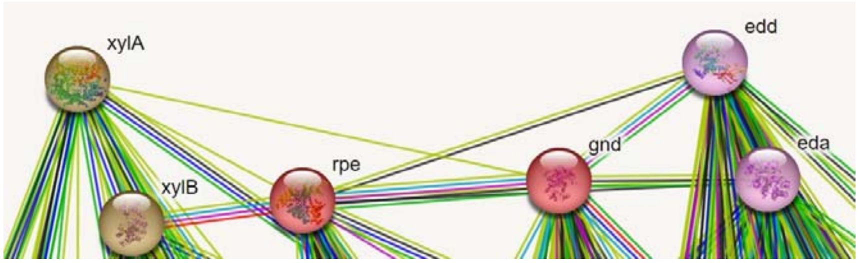


**Figure S6. Pathway analysis using Cytoscape StringApp software. a-d** Separate networks for xylose conversion, glycolysis, pentose-phosphate, and all proteins related to $CO_2$ emission/fixation. **e,f** Merged networks to identify the enzyme most closely associated with the xylose conversion pathway and possessing $CO_2$ fixation capability.

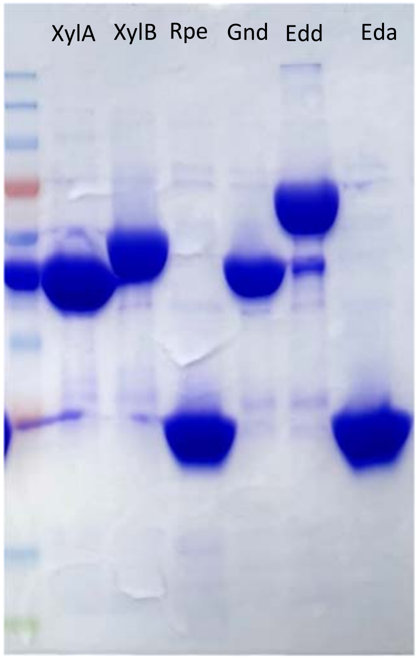


**Figure S7**. **SDS-PAGE of the six enzymes immobilized within the chips.** The six His-tagged enzymes expressed in *E. coli* were purified using HisPur™ Ni-NTA Resin. 5 µl of the purified proteins were mixed with 4X Laemmli sample buffer and loaded in a 12% Mini-PROTEAN® TGX™ Precast Protein Gel. SDS-PAGE was performed at 145 V for 50 min and the resulting gel was stained with Coomassie Brilliant Blue R-250.

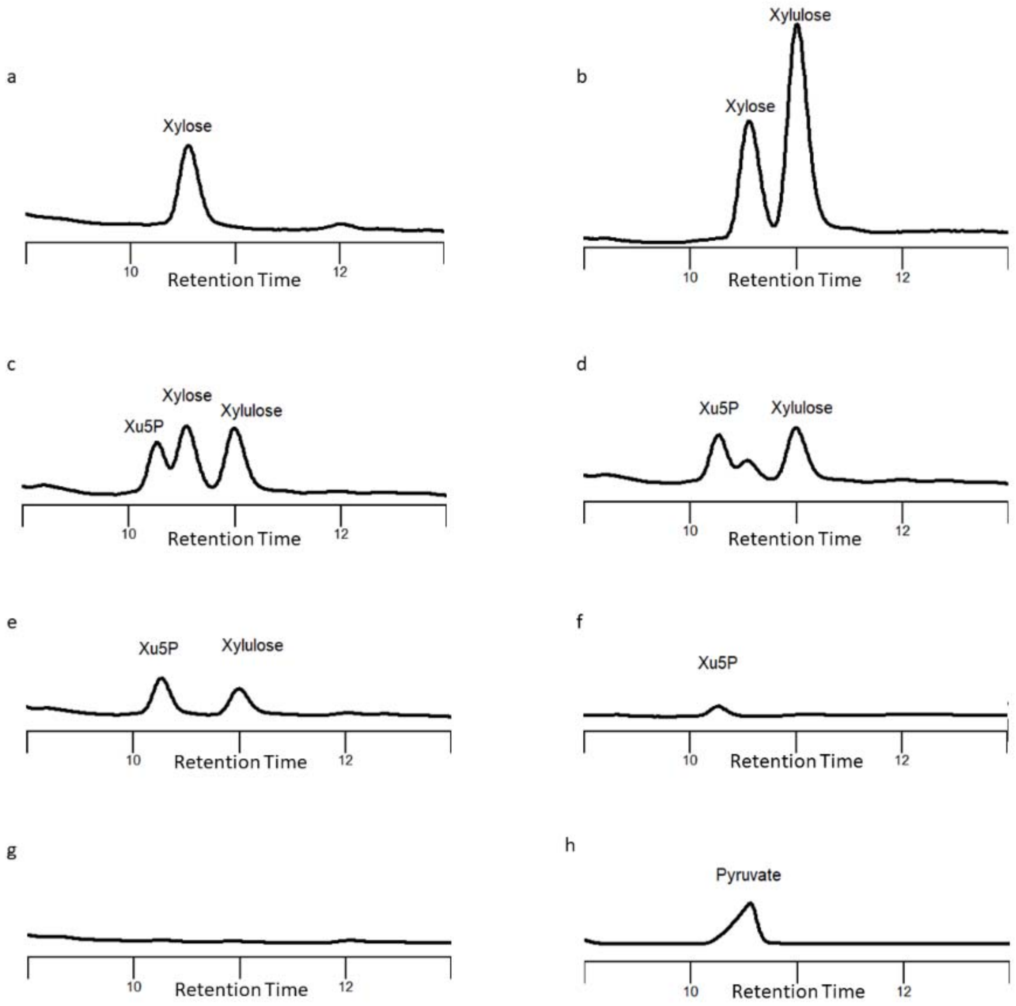


**Figure S8. HPLC chromatogram. a.** Xylose standard retention time. **b.** xylose conversion to xylulose by XylA enzyme. **c.** Xylose conversion to xylulose and xylulose 5-phosphate (Xu5P) through the XylA and XylB enzymes reactions. **d-g.** Progression of xylose, xylulose, and Xu5P consumption by subsequent enzymes in the pathway without Eda over a 24-hour period. **h.** Pyruvate production in the pathway accompanied by Eda enzyme over 24-hour reactions. Pyruvate exhibits a retention time similar to xylose, but its absorbance at 300 nm allows differentiation from xylose.

**Table S1.** Primers used for plasmid construction.

| Primer | Sequence |
|---|---|
| **FxylA** | CATCACCATCATCACCACAGCCAGGATCCGCAAGCCTATTTTGACCAGCTCGAT |
| **RxylA** | GCGGTTTCTTTACCAGACTCGAGTTATTTGTCGAACAGATAATGGTTTACC |
| **FxylB** | CATCACCATCATCACCACAGCCAGGATCCGTATATCGGGATAGATCTTGGCACCTC |
| **RxylB** | GCGGTTTCTTTACCAGACTCGAGTTACGCCATTAATGGCAGAAGTTG |
| **Frpe** | CATCACCATCATCACCACAGCCAGGATCCGAAACAGTATTTGATTGCCCCCTCA |
| **Rrpe** | GCGGTTTCTTTACCAGACTCGAGTTATTCATGACTTACCTTTGCCAGTTCA |
| **Fgnd** | CATCACCATCATCACCACAGCCAGGATCCGTCCAAGCAACAGATCGGCGTAGTC |
| **Rgnd** | GCGGTTTCTTTACCAGACTCGAGTTAATCCAGCCATTCGGTATGGAACA |
| **Fedd** | CATCACCATCATCACCACAGCCAGGATCCGAATCCACAATTGTTACGCGTAACAAATC |
| **Redd** | GCGGTTTCTTTACCAGACTCGAGTTAAAAAGTGATACAGGTTGCGCCC |
| **Feda** | CATCACCATCATCACCACAGCCAGGATCCGAAAAACTGGAAAACAAGTGCAGAATCAA |
| **Reda** | GCGGTTTCTTTACCAGACTCGAGTTACAGCTTAGCGCCTTCTACAGCTTC |
| **Fpet** | CGATGCGTCCGGCGTAGAGG |
| **Rpet** | GCTAGTTATTGCTCAGCGGTGGC |
